# Mechanistic Modeling of Biochemical Systems Without A Priori Parameter Values Using the Design Space Toolbox v.3.0

**DOI:** 10.1101/2020.01.30.927657

**Authors:** Miguel Á. Valderrama-Gómez, Jason G. Lomnitz, Rick A. Fasani, Michael A. Savageau

## Abstract

Mechanistic models of biochemical systems provide a rigorous kinetics-based description of various biological phenomena. They are indispensable to elucidate biological design principles and to devise and engineer systems with novel functionalities. To date, mathematical analysis and characterization of these models remain a challenging endeavor, the main difficulty being the lack of information for most system parameters. Here, we introduce the Design Space Toolbox v.3.0 (DST3), a software implementation of the Design Space formalism that enables mechanistic modeling of complex biological processes without requiring previous knowledge of the parameter values involved. This is achieved by making use of a phenotype-centric modeling approach, in which the system is first decomposed into a series of biochemical phenotypes. Parameter values realizing phenotypes of interest are predicted in a second step. DST3 represents the most generally applicable implementation of the Design Space formalism to date and offers unique advantages over earlier versions. By expanding the capabilities of the Design Space formalism and streamlining its distribution, DST3 represents a valuable tool for elucidating biological design principles and guiding the design and optimization of novel synthetic circuits.

## 1. Introduction

Mechanistic models have the advantage of being biologically realistic and the potential to rigorously define and predict biochemical phenotypes. However, the vast number of kinetic parameters whose values are largely unknown is a bottleneck limiting their use. Thus, current approaches to determining the phenotype focus first on <u>estimating</u> parameter values for the underlying biochemistry, typically through a mixture of ad-hoc experimentation and computationally inefficient high-dimensional numerical search, and then exploring the model’s repertoire by <u>simulation</u>. While these strategies have been used to fully characterize small systems in the pre-genomic era, a mechanistic understanding of systems, even of moderate size, derived from genotype data remains elusive.

Here, we describe computational tools for the implementation of an alternative post-genomic approach –the Design Space Analysis, that first *<u>analytically</u>* determines the space of possible phenotypes for a given system architecture (which can be inferred from high-throughput data) and then *<u>predicts</u>* parameter values for their realization, predictions that can guide experimentation and further numerical analysis.

The theoretical foundation of the Design Space formalism was laid back in the 70’s (Savageau 1969, Savageau 1971a, Savageau 1971b, Savageau 1979). This early work introduced the concept of S-systems and their mathematical characterization regarding dynamic stability of steady states, logarithmic gains for signal amplification, and parameter sensitivities for robustness. Recently, these concepts were integrated to describe a generic approach to the construction of the *Design Space*, a structured parameter space in which qualitatively distinct biochemical phenotypes can be identified, counted and located (Savageau et al 2009). Linking regions of the parameter space with biochemical phenotypes has allowed the elucidation of design principles and the introduction of a radically new phenotype-centric modeling strategy (Lomnitz and Savageau 2016b, Valderrama-Gomez and Savageau 2018). Over the last decade, two computational implementations of the Design Space formalism have been developed. Rick Fasani first introduced the Design Space Toolbox for MATLAB (DST1), a formal software implementation automating key steps of this methodology. Fasani’s contribution included an elegant mathematical description of the Design Space and a detailed explanation of its construction (Fasani and Savageau, 2010). Later, Jason Lomnitz introduced the Design Space Toolbox V2 (DST2) (Lomnitz and Savageau, 2016b). DST2 consisted of a collection of tools comprised of a stand-alone library, written in the C language, that implements its own symbolic algebra engine and leverages open-source compiled libraries for linear algebra and linear optimization (via the GLPK library). By using multi-threaded concurrent algorithms to speed up calculations, DST2 took advantage of the parallelizable nature of the Design Space approach by analyzing each biochemical phenotype of the system independently.

Here, we introduce the Design Space Toolbox v.3.0 (DST3). This new version builds on DST2 to expand the capabilities of the Design Space formalism by allowing the automatic identification and mathematical characterization of additional phenotypes arising from critically important under-determined cases. These special cases emerge from cycles, metabolic imbalances and conservation constraints present in many biochemical systems. The expanded computational engine of DST3, its C-library, is now able to handle these singularities when they appear individually and simultaneously. Keeping users with limited programming experience in mind, DST3 offers an improved and more stable IPython-based user interface. New functionalities allow, among other things, calculation of the product of the tolerances for all parameters in log-coordinates, a proxy for a phenotype’s volume in parameter space and for its associated global robustness. Additionally, solvers for systems of both ordinary differential (ODE) and differential algebraic (DAE) equations were incorporated, thus allowing a fully integrated numerical characterization of the *Full System*. The analysis of the full system provides a means to assess the accuracy of predictions made by DST3, which are based on underlying S-Systems.

This integrated suite of computational algorithms for the efficient prediction of parameter values and analysis of the phenotypic repertoire is provided in a user-focused environment for navigating the resulting space of phenotypes and identifying biologically relevant design principles. These innovations will facilitate deterministic and stochastic simulations that require parameter values, will accelerate both hypothesis discrimination in systems biology and the design cycle in synthetic biology, and will enable investigators to achieve predictive understanding of biomolecular phenotypes from genotype.

In order to simplify the installation process and to guarantee usage across different operating systems – Widows, macOS and Linux – we generated a Docker Image for DST3. This effectively renders Docker the only software dependency of DST3. Altogether, we believe that innovations contained in DST3 will greatly boost the application of the Design Space formalism for the analysis of biochemical systems and the elucidation of their underlying design principles.

This manuscript is divided into three main sections. The first section reviews briefly key concepts of the Design Space formalism (see Savageau and Fasani, 2009 and Fasani and Savageau, 2010 for a more detailed theoretical treatment) needed to understand the advances described in this work. In the second section we build on these concepts to develop mathematical strategies aimed at resolving matrix singularities arising from system topologies containing cycles, moiety conservations, and metabolic imbalances leading to blow-ups/-downs. In the third section, we illustrate the capabilities of DST3 by analyzing a case study of a biochemical system exhibiting multiple, nested singularities. The Methods section provides details on the software architecture of DST3, the different ways to access its computational capabilities, and the installation instructions via Docker.

## 2. Review of Key Concepts

Biochemical systems described by the power-law functions of chemical kinetics and the rational functions of biochemical kinetics can be represented by generalized mass action (GMA) kinetics (Savageau and Voit, 1987) of the form:

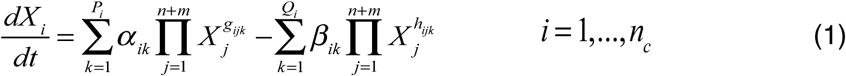

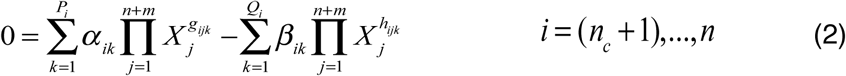

where *X*_*i*_ represents the concentration of a chemical species of interest in a system containing a total of *n* dependent and *m* independent variables. In general, dependent variables can be split into two groups: chemical variables, for which a differential equation exists, and auxiliary variables, for which algebraic constraints are defined. Each group contains *n*_*c*_ and *n* − *n*_*c*_ members, respectively. *α*_*ik*_ and *β*_*ik*_ represent rate constants, while *g*_*ijk*_ and *h*_*ijk*_ are kinetic orders. *P*_*i*_ and *Q*_*i*_ are the number of positive and negative terms in the *i*-th equation, respectively.

Variables for which a differential equation or algebraic constraint are not defined are treated as parameters. For any system in steady state, one of the *P* positive terms and one of the *Q* negative terms will dominate over the others in each one of the *n* equations in the system. This gives rise to a so-called *dominant S-System*, which can be generically described by Eq. 3:

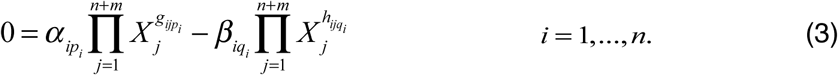

With *p*_*i*_ and *q*_*i*_ being the indices of the dominant positive and dominant negative term in the *i*-th equation, respectively. The validity of the dominant S-System implies certain conditions, which are represented by inequalities of the form

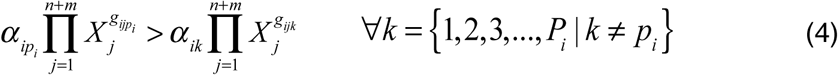

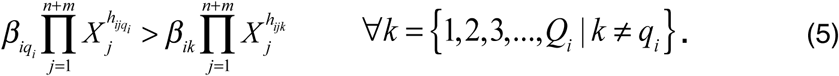

Here, *k* represents indices of corresponding non-dominant terms. Steady state concentrations of the dependent variables can be obtained in three steps. By rearranging Eq. 3 and taking logarithms, one obtains Eq. 6:

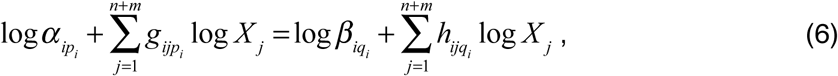

which can be written in matrix form as:

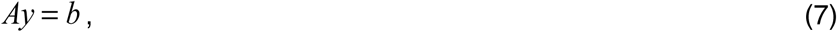

where 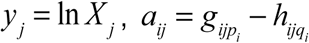 and *b*_*i*_ = ln(*β*_*in*_ / *α*_*iq*_). In a second step, dependent (*y*_*D*_) and independent (*y*_*I*_) variables are split to obtain:

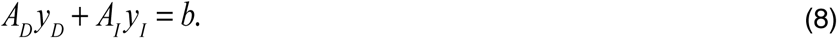

The vector of dependent concentration variables *y*_*D*_ can be obtained in a third step by matrix operations:

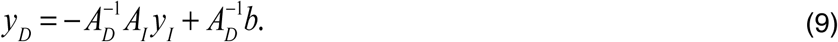

The vector of dependent flux variables log*V*_*i*_ is obtained by matrix multiplication:

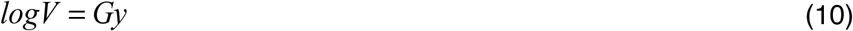

## 3. Strategies for Treating Three Types of Singularities

Earlier versions of the Design Space Toolbox (DST1 and DST2) have exclusively dealt with cases for which the inverse of the matrix *A*_*D*_ exists. However, the matrix *A*_*D*_ becomes singular for certain system topologies, which have previously been dealt with on a case-by-case basis. The theory behind these cases is well-known, and here we have implemented the automatic handling of these cases. The strategies are illustrated by means of simple examples treated briefly below with mathematical details given in the Supplemental Information.

### 3.1 Cycles are resolved by considering global dominance equations

Cycles of reactions are a common feature of biochemical systems. They typically have a number of input and output fluxes. A simple example is the FNR global regulator of *Escherichia coli* that exists in a cycle with three forms having one influx and two effluxes (Tolla et al. 2015). Consider the simple system shown in Fig 1A., in which species *X*_1_, *X*_2_ and *X*_3_ interact to form a substrate cycle driven far from thermodynamic equilibrium. This example is deliberately selected to focus on the mathematical details of the singularity contained in the kinetic equations and on the strategy that resolves it. We start by setting up equations to describe the change in the concentration of each chemical species over time. Mass action kinetics are used to generate the rate laws describing the flux through each reaction. The resulting expressions are then combined by means of Kirchhoff’s node law to generate balance equations for each metabolite in the system.

**Figure 1.**
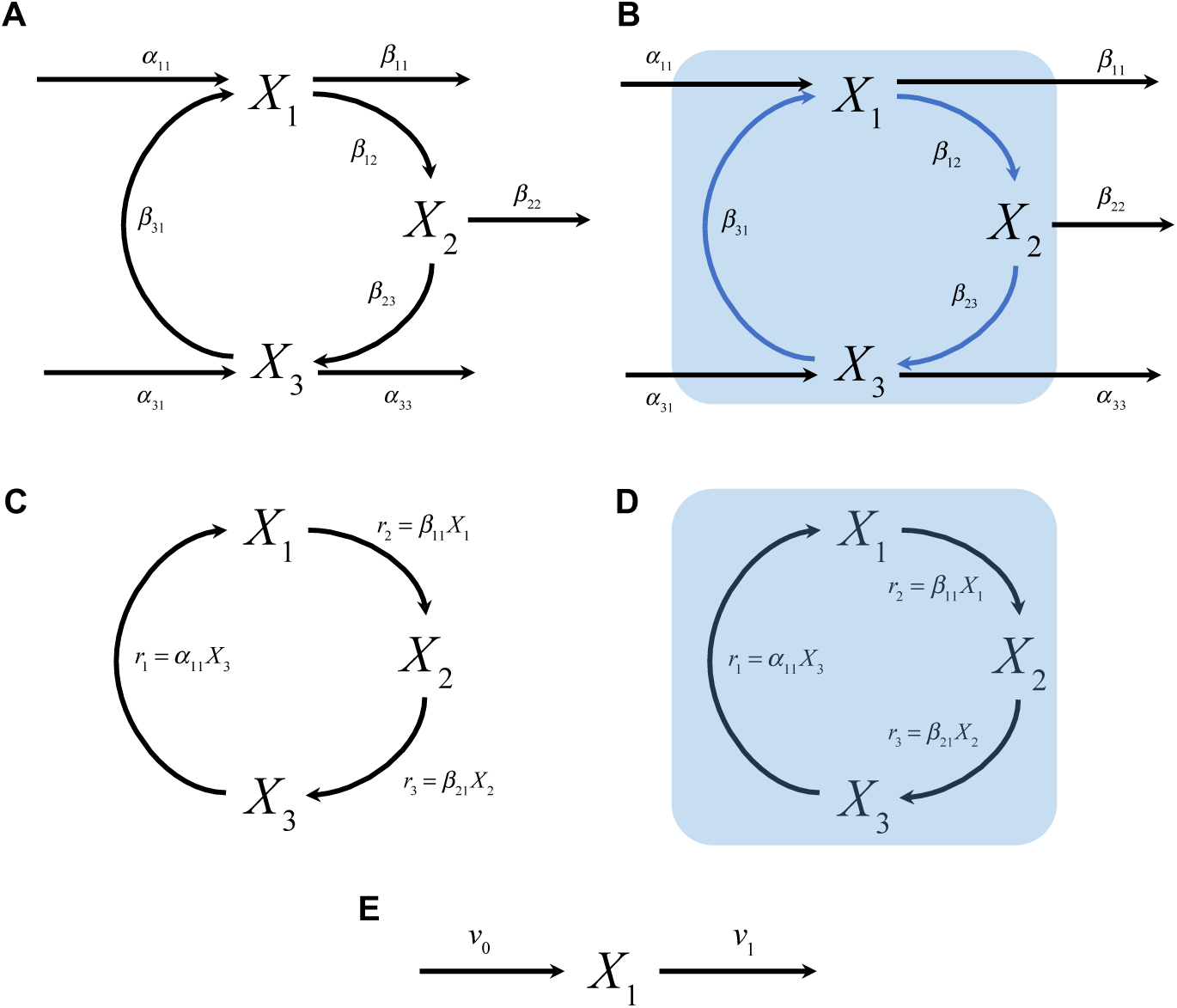
Systems Containing Cycles, Conservation Constraints and Metabolic Imbalances. **A)** A cycle comprised of three metabolic species is shown. Mass action kinetics is used to mathematically describe the flux through each reaction. Rate constants for each one of these reactions are shown. **B)** If the fluxes represented by the blue arrows dominate over the others, a singularity arises that prevents the system from having a unique steady state solution. This singularity is resolved by introducing a mass balance equation around a control volume (blue rectangle) containing the cycle. **C)** Three-component system containing one conservation constraint. Fluxes to and from metabolite pools *X*_1_, *X*_2_ and *X*_3_ are mathematically described using mass action kinetics. **D)** No fluxes enter or leave the control volume (blue rectangle) around metabolites involved in the conservation. **E)** Metabolic pool with one input and one output flux. Depending on the numerical values of fluxes *v*_0_ and *v*_1_, the concentration of *X*_1_ can steadily increase (*v*_0_ > *v*_1_), decrease (*v*_0_ < *v*_1_) or remain unchanged over time (*v*_0_ = *v*_1_).

The Design Space formalism can be applied to decompose this set of equations into different cases, each having a unique set of dominant terms and being valid within a specific region in parameter space. One such case for this system is case number 27, with case signature [22 11 21]. This signature contains three pairs of indices, one for each equation, indicating the identity of the positive and negative term dominating in each equation. Visual inspection of the matrix *A*_*D*_ for this case reveals the presence of a linear dependency among its rows, and, thus, there is no unique steady state solution. Nevertheless, the system of algebraic equations is consistent and a solution (or set of solutions) can be found by analyzing global dominance conditions on the influxes and effuxes that describe a mass balance around the cycle present in this system.

Once the extended sub-system has been automatically set up, the Design Space formalism can be applied to identify valid sub-cases that resolve the cyclical case. Table S1 shows S-system equations for each one of the six valid cases generated from the extended sub-system. Note the special form of these equations. S-systems originating from the dominance analysis of the global dominance equation are used to replace the differential equation for the pool with the dominant efflux. For instance, the expression *α*_11_ − *β*_11_*X*_1_ (obtained when the first positive and first negative term in the global dominance equation are dominant) is used to replace the differential equation for *X*_1_ (refer to sub-case 1 in Table S1). Additionally, this expression is scaled to match the coefficient of the negative term in the full system (i.e., the original set of equations). Consider for instance the expression *α*_11_ − 2*β*_33_*X*_3_, which is obtained when the first positive and third negative term in the global dominance equation are dominant. Since the coefficient of the negative term in the original equation is 1, the scaled expression 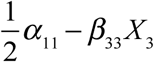 is used to construct sub-case 3 of Table S1.

Due to the special way in which terms stemming from global dominance are used to construct equations for sub-cases, a three-digit case signature is introduced. This allows for tracking the origin of terms that make up the S-systems of these sub-cases. In addition to the indices of dominant positive and negative terms contained in the traditional two-digit signature, the extended three-digit signature contains the index of the equation from which its positive dominant term originated. Consider for instance the case signature for sub-case 27_5 in Table S1 and its associated S-system. The signature [22 **3**12 21] dictates that differential equations for pools *X*_1_ and *X*_3_ are constructed by picking dominant terms in the traditional way, while the differential equation for pool *X*_2_ is made from the first positive term of the *third* equation and the second negative term of the *second* equation.

### 3.2 Conserved moieties are handled by considering the total size of conserved pools

Pools of metabolites with constant total concentration, on some time scale, are a common feature of complex metabolic and signaling systems. They involve conserved moieties, which are groups of atoms that remain intact in all reactions of a system. AMP, NAD^+^, and NADP^+^ are prominent examples of conserved moieties in energy metabolism (Haraldsdóttir and Fleming, 2016). Consider the simple system in Fig. 1C with three components linked by a conservation relationship. As in the case of the cycles in *Section 3.1*, the presence of conservations would cause the matrix *A*_*D*_ to be singular, since the three concentrations are not independent. Nevertheless, steady state solutions can be obtained by discarding one of the differential equations and adding the algebraic constraint that the sum of the three concentrations must equal their conserved amount.

The analysis of the differential-algebraic system using the Design Space formalism involves the usual generation of cases by picking dominant terms for each of the equations of the system. The three cases that result are shown in Table S2. Note that each case is defined by only two differential equations and one algebraic constraint. In order to capture the special way in which the equations are constructed for each case, the indices of the differential equation being deleted are set to zero in the case signature. Case 3 for instance, in which differential equation for pool *X*_3_ is missing, has a case signature of [11 11 **00** 13] to reflect this fact.

### 3.3 Metabolic Imbalances are treated by considering knife-edge conditions

Imbalances are frequently encountered in the metabolism of engineered microbial strains (Dahl et al. 2013, George et al 2014, Alonso-Gutierrez et al. 2017) and in inborn metabolic diseases such as phenylketonuria (Levy 1999) and maple syrup urine disease (Haymond et al. 1973), often by the excretion of some metabolite. Consider the simplest example of a metabolic pool as shown in Fig. 1E with one input (*v*_0_) and one output (*v*_1_) flux, where *v*_0_ is a constant, and the output flux is described by a Michaelis-Menten rate law 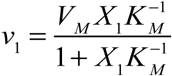. *K*_*M*_ and *V*_*M*_ represent the Michaelis constant and maximal reaction rate, respectively. The change in concentration of metabolite *X*_1_ over time can be described by the generalized mass action system

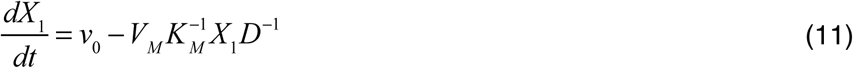

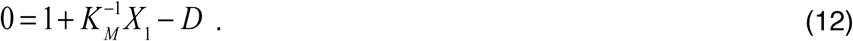

where *D* represents an auxiliary variable introduced in the recasting process to describe the denominator of the Michaelis-Menten rate law. Let us now consider the equations for the case with signature [11 21]

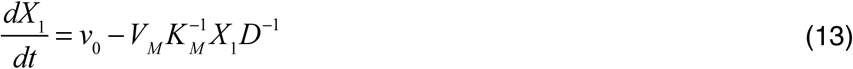

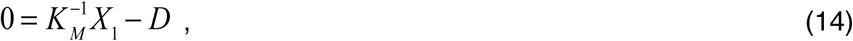

which together with its associated dominance condition:

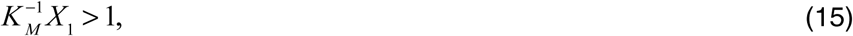

imply:

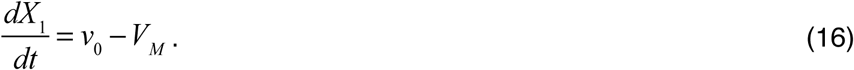

This system does not have a steady state solution. Indeed, Eq. 16 only provides a consistency condition for the concentration of *X*_1_ to remain unchanged over time: 0 = *v*_0_ − *V*_*M*_. We will refer to this kind of constraint as a *knife-edge condition*. In general, we are interested in the behavior of the system when knife-edge conditions are not satisfied, i.e., *v*_0_ ≠ *V*_*M*_. Violating the knife-edge condition in a specific direction implies an extreme value for *X*_1_ : *X*_1_ → ∞ or *X*_1_ → 0. The validity of either situation is assessed by checking the validity of the associated dominance conditions, as shown in Table S3. Taken together, these results indicate that for the system shown in Fig. 1E, the concentration of the pool *X*_1_ will steadily increase over time, i.e., it will *blow up* if the system’s parameters fulfill the conditions 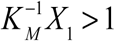 and 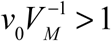.

Note that the case with signature [11 11], and associated dominance condition, has a conventional steady state solution given by 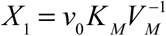 for 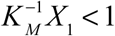. The general procedure for treating cases with multiple knife-edge conditions is presented in the Supplemental Information.

## 4. Analysis of a Biochemical System Exhibiting Multiple Singularities

For a case study, we have selected a model system that is important in metabolic engineering for the detoxification of environmental pollutants and well as the production of high-quality chemical precursors. There are many microbes capable of these industrially important processes in the context of toxic environmental hydrocarbons. Although they may have similar if not identical pathways for these functions, their genomic architectures exhibit major differences that are not well understood (Jiménez et al. 2002, Harwood and Parales 1996). One of the simplest of these architectures for the transport and catabolism of protocatechuate is found in *Acinetobacter* sp. strain ADP1. Its genetic system encodes a single polycistronic mRNA for transporter and catabolic enzymes of the protocatechuate specific pathway, as well as several shared enzymes (Trautwein and Gerischer 2001, Dal et al. 2005). For our purposes, we shall focus only on the protocatechuate specific pathway illustrated in Figure 2A. The system is composed of a signaling cascade, a gene circuit and a metabolic module. The transcription factor (PcaU) functions as both a repressor, *U*_1_, and an activator, *U*_2_, and has a conserved total concentration of *U*_*T*_ = *U*_1_ + *U*_2_. It controls synthesis of a polycistronic mRNA, *M*_3_, that encodes the transporter (PcaK), *T*_4_, and several enzymes, three of which constitute this pathway: protocatechuate 3,4-dioxygenase (PcaGH), *GH*_5_, *β* -carboxy-*cis,cis*-muconate cycloisomerase (PcaB), *B*_6_, and 4-carboxymuconolactone decarboxylase (PcaC), *C*_7_. The environmentally supplied substrate protocatechuate, protocatechuate, *P*_0_, is transported into the cell where it becomes intracellular protocatechuate, *P*_8_, which is both a metabolic intermediate and the natural inducer for the transcription factor. The following metabolites in the pathway are β-carboxy-*cis,cis*-muconate, *CM*_9_, and γ-carboxymuconolactone, *CL*_10_ (Dal et al. 2005).

**Figure 2.**
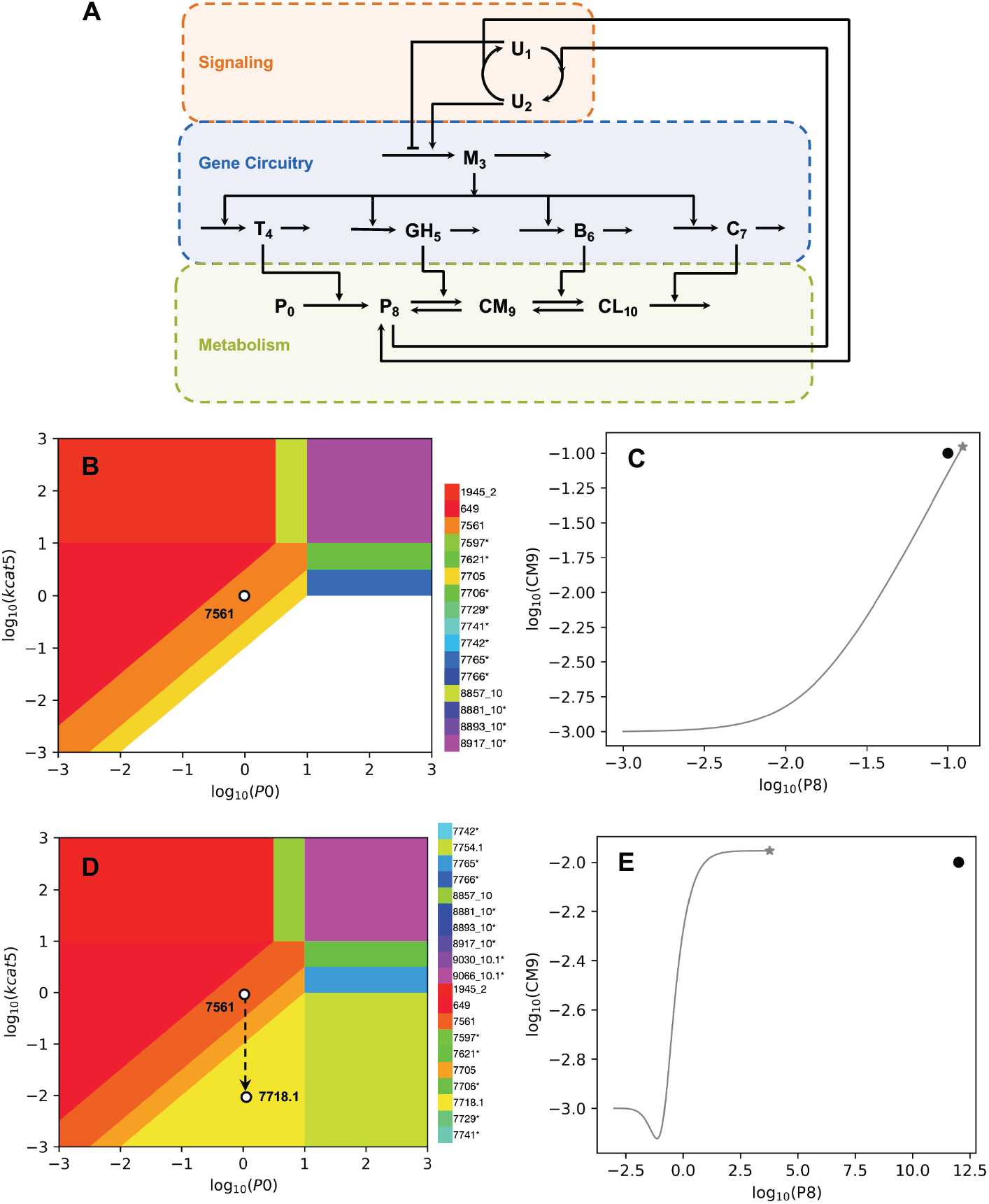
Integrated System Exhibiting Multiple Singularities and its Characterization by DST3. **A)** The signaling module processes an inducer signal *P*_*8*_, which stimulates the conversion of the transcription factor from the repressor *U*_*1*_ into the activator *U*_*2*_ form. The regulator controls the production of a polycistronic mRNA molecule *M*_*3*_, from which four proteins *T*_*4*_, *GH*_5_, *B*_6_ and *C*_7_ are translated. Transporter *T*_*4*_ catalyzes the import of metabolite *P*_8_ into the cell from an external pool *P*_0_. Enzymes *GH*_5_ and *B*_6_ then catalyze the reversible conversion of *P*_8_ into *CM*_9_ and *CM*_9_ into *CL*_10_. The last enzyme *C*_7_ catalyzes the conversion of *CL*_10_ into the end-product β-ketoadipate enol-lactone (not shown). **B)** A **Design Space plot** around phenotype **7561** is shown. The white dot represents the operating point of the system, which is automatically calculated using the **Analyze Case** tab of the DST3 user interface. Parameter values for this operating point are: K_1_ = 10, K_2_ = 1.0, K_M4_ = 10.0, K_M5f_ = 1.0, K_M5r_ = 1.0, K_M6f_ = 1.0, K_M6r_ = 1.0, K_M7_ = 1.0, K_eq5_ = 1.0, K_eq6_ = 1.0, U_T_ = 3.16, P_0_ = 1.0, α_1_ = 1.0, α_3basal_ = 0.01, α_3max_ = 1.0, α_3min_ = 1.0, α_4_ = 1.0, α_5_ = 1.0, α_6_ = 1.0, α_7_ = 10.0, *β*_1_ = 1.0, *β*_3_ = 1.0, *β*_4_ = 1.0, *β*_5_ = 1.0, *β*_6_ = 1.0, *β*_7_ = 1.0, k_cat4_ = 1.0, k_cat5_ = 1.0, k_cat6_ = 1.0, k_cat7_ = 1.0; Kinetic order(s): m = 2, p = 2; Parametric constraints: α_3max_ > α_3basal_ > α_3min_. **C)** A **trajectories plot** is used to characterize the operating point shown in panel **B**. The solid grey line represents the evolution of the system starting from an initial condition in which all concentrations are set to 0.001. The grey star represents the steady state reached by the system after numerical integration for 500 time units. The black dot next to the grey star represents the steady state predicted by DST3 for phenotype 7561 using linear algebra. **D)** The operating point of the system has been modified by decreasing k_cat5_ from 1.0 to 0.01 so that it is now contained within the region of blow-up phenotype **7718.1**. The steady state of the new operating point of the system is characterized using a **trajectories plot** (panel **E**), in which the temporal behavior of metabolite pools *P*_8_ and *CM*_9_ is shown. *CM*_9_ quickly reaches a steady state of approximately 0.01, while the concentration of *P*_8_ continuously increases over time and does not reach a steady state. The grey star represents the state of the system after numerical integration for 50000 time units. When integrated for a longer time period, the grey star would move further to the right at a constant **CM**_**9**_ concentration, till it eventually reaches and passes the location of the black dot. Initial conditions are the same as in panel **C**.

This system contains multiple singularities. Conservation relationships from the transcription factor, cycles from the reversible enzymatic reactions within the metabolic pathway, and blow-ups from imbalances within the same pathway. The dynamics of the system can be described by a set of differential algebraic equations involving 30 unknown parameter values. Refer to Eqs. S34 – S45 in the Supplemental Information for details.

### 4.1 Filtering the phenotypic repertoire for phenotypes of interest

Enumerating the phenotypic repertoire of a system is typically the first step in the phenotype-centric modeling strategy. Even systems of moderate size can exhibit a surprisingly large number of biochemical phenotypes. Therefore, the second important step is to filter the repertoire for the phenotype of interest. For example, filtering for cases with 2 eigenvalues with positive real part can be used to identify oscillatory phenotypes (Lomnitz and Savageau, 2014), filtering for cases with 1 eigenvalue with positive real part can be used to identify multi-stability and hysteresis (Fasani and Savageau, 2013), and filtering for a logical function consisting of a pattern of dependent variables that increase, decrease or remain unchanged in response to a change in an independent variable can be used for model discrimination (Lomnitz and Savageau, 2016a). Since all of the phenotype characteristics can be exported from DST3 to an Excel spread sheet, and this allows for many types of user-defined filters that can be customized to meet the user’s needs (see part 3 of the tutorial contained within the DST3 Docker image for an example).

Here, we show how one can progressively filter the repertoire of the protocatechuate system to narrow the focus on phenotypes of interest. If we allow for all possibilities, the DST3 shows that the system represented in Fig. 2A is capable of exhibiting 3722 phenotypes. However, we can progressively filter this list automatically to include only those phenotypes of interest. In the context of this case study, we are interested in the steady states of this system that maximizes the pathway flux, while minimizing the accumulation of toxic intermediates. First, if we filter for phenotypes that are non-pathological by not checking for blow-ups, i.e., that do not have imbalances resulting in concentrations that continuously increase or decrease, then the number of non-pathological phenotypes is 384 and they are all stable (all eigenvalues have a negative real part). Second, if we filter these for phenotypes that respond to changes in the environmental substrate **P**_**0**_, by requiring a non-zero logarithmic gain in metabolite concentrations in response to a change in substrate, then there are only 192 responders. Third, if we filter these for phenotypes that are inducible, by requiring a non-zero logarithmic gain in mRNA in response to changes in substrate, then there are only 64 inducible responders. Finally, if we filter the inducible phenotypes for specific non-zero logarithmic gains in mRNA, then we find only three values: L(**M**_**3**_,**P**_**0**_)=2 with 32 examples, L(**M**_**3**_,**P**_**0**_)=4 with 28 examples, and L(**M**_**3**_,**P**_**0**_)=6 with 4 examples.

Following this initial screening, the DST3 can be used to characterize automatically the inducible responders by comparing them on the basis of three functional criteria: Global robustness to a change in phenotype, energy index (maximum flux with minimum production of protein machinery), and toxicity index (maximum flux with minimum accumulation of toxic intermediates). Global robustness is determined by the product of the global tolerances for all of the parameters of the system, which is a proxy for the volume of the phenotype’s polytope in the system Design Space. We define the energy index as the cost/benefit determined by the ratio of the logarithmic gain in mRNA, which is a proxy for the increased expenditure of energy for protein production, to the logarithmic gain in the pathway flux produced, *energy index* = *L*(*M* _3_, *P*_0_) / *L*(*F, P*_0_). The toxicity index is the cost/benefit determined by the ratio of the logarithmic gain in the toxic intermediate, protocatechuate (*P*_8_), to the logarithmic gain in the pathway flux produced, *toxicity index* = *L*(*P*_8_, *P*_0_) / *L*(*F, P*_0_).

The results summarized in Table 1 show that 32 of the 64 phenotypes have the best global robustness, best energy index, and best toxicity index. The next 28 phenotypes have intermediate values for these three criteria and the remaining 4 phenotypes have the worst global robustness, and worst energy index, and the worst toxicity index. There is a clear trade-off revealed by this analysis. The pathway flux can be increased by moving from the phenotypes in the first group to those in the third group, but only by sacrificing global robustness, energy efficiency, and toxicity.

**Table 1.** *Repertoire of 64 phenotypes that are responding to substrate, are stable and inducible.* This table was generated for the system described by Eqs. S46 – S62 of the Supplemental Information using the tab **Phenotypic Repertoire** of the **Main Menu** of the user interface of DST3 and filtered for desirable phenotypes as described in the main text.

|  | Group |  |  |
| --- | --- | --- | --- |
|  | 1 | 2 | 3 |
| Number of phenotypes in group | 32 | 28 | 4 |
| Representative phenotype | 7561 | 7657 | 7665 |
| $L(\mathbf{M}_3, \mathbf{P}_0)$ | 2 | 4 | 6 |
| $L(\mathbf{P}_8, \mathbf{P}_0)$ | 1 | 2 | 3 |
| $L(\mathbf{CM}_9, \mathbf{P}_0)$ | 1 | 1 | 2 |
| $L(\mathbf{CL}_{10}, \mathbf{P}_0)$ | 1 | 1 | 1 |
| $L(\mathbf{F}, \mathbf{P}_0)$ | 3 | 5 | 7 |
| Normalized volume of representative phenotype | $1.94 \times 10^{-08}$ | $7.57 \times 10^{-12}$ | $9.72 \times 10^{-16}$ |
| Total normalized volume of group | $5.61 \times 10^{-7}$ | $6.32 \times 10^{-11}$ | $3.40 \times 10^{-15}$ |
| Energy index | 0.67 | 0.80 | 0.86 |
| Toxicity index | 0.33 | 0.40 | 0.43 |

### 4.2 Phenotype analysis reveals dynamic properties of the system

Consider phenotype number **7561** as a representative of the best class, which could conceivably have been selected in nature. Since we have the ability to characterize the full system exhibiting this phenotype, we can ask if there might be a strategy for further improving its performance or for avoiding dysfunction through rational engineering. We start by using DST3 to predict specific values for each one of the 30 parameters required to fully define an operating point for this phenotype of the system. This is done using the **Analyze Case** tab within the **Main Menu** of the DST3 user interface. We opt to locate the operating point of the system within phenotype **7561**, but an analogous analysis can be performed for any other phenotype. An obvious dysfunction occurs when there is a violation of one of the most basic design principles; namely, the maximal velocity of a downstream enzyme should be greater than that of the upstream enzymes in the pathway (Savageau et al, 2009). The relation of this pathology to the phenotype **7561** is made evident in a **Design Space plot** with the turnover number for the enzyme converting metabolite *P*_*8*_ to *CM*_*9*_, *k*_*cat5*_, on the y-axis. Fig. 2B shows the location of the operating point as a white dot, which is contained, as desired, within phenotype **7561** (orange polytope). In this view of the system Design Space we have only included the physiologically relevant phenotypes; the white blank region indicates the location of pathological phenotypes with a blowup. The dynamical behavior of the full system can be studied using a **trajectories plot**. This plot shows the evolution of the system starting from its initial condition to reach its steady state operating point, which is marked by a grey star in Fig. 2C. The black dot in this figure represents the steady state prediction made by DST3 for phenotype **7561**. The relative position of the black dot and the gray star in the trajectories plot demonstrates the accuracy of DST3 when approximating steady states. Additionally, as expected from the number of positive eigenvalues for phenotype **7561**, the full system exhibits a single, stable steady state.

It is also possible to identify the nature of the pathological phenotypes. By clicking the “Check for Blowups” option in the construction of the Design Space, we obtain Fig. 2D, which has four additional phenotypes displayed (7718.1, 7754.1, 9030_10.1 and 9066_10.1). The dynamical nature of blowing phenotypes contained in the lower portion of the **Design Space plot** can be trivially predicted. Decreasing the numerical value of *k*_*cat5*_ while keeping all other parameter values constant at the operating point of the system leads to a continual increase of the toxic intermediate *P*_*8*_ while the intermediate *CM*_9_ quickly reaches its steady state value, as shown in Fig 2E. In summary, a comparison of the results obtained by numerical integration of the full system (Fig. 2C and E) demonstrates the ability of DST3 to predict steady state values when they exist and predict the blowing nature of a variable when it does not have a steady state solution.

### 4.3 Logarithmic gains can guide the design of engineering strategies

Once a stable and globally robust operating point for a given system has been identified, one might be interested in finding strategies to increase the flux through a specific metabolic pathway or to increase the steady state concentration of certain intermediate metabolites. Here, we exemplify how an analysis of logarithmic gains can be used to identify such strategies. *Logarithmic gains* are amplification factors relating changes in input signals, independent variables, to the resulting changes in output signals (dependent variables). The term *parameter sensitivity* is used instead of logarithmic gain when the effect of varying a parameter on a dependent variable is analyzed. These parameter (in)sensitivities represent the local robustness of a system, in contrast to the global robustness provided by the volume of a phenotype in Design Space. Both logarithmic gains and parameter sensitivities are properties that depend exclusively on the kinetic orders of the system and can be calculated for concentrations or fluxes (Savageau 1971a). DST3 allows the calculation of logarithmic gains and parameter sensitivities using the tab **Analyze Case** of the **Main Menu** in the user interface. For simplicity, we will use the term logarithmic gain for both logarithmic gains and parameter sensitivities. Table 2 lists logarithmic gains for the phenotype **7561**, the representative phenotype of the first group of phenotypes with desired properties (Table 1).

**Table 2.** *Logarithmic Gains for Phenotype* ***7561***. This table was generated using the tab **Analyze Case** of the **Main Menu** in the DST3 user interface. Shown are logarithmic gains for the steady state concentration of the mRNA molecule **M**_**3**_, the three pathway intermediates **P**_**8**_, **CM**_**9**_ and **CL**_**10**_, and the flux through the metabolic pathway **F**. Parameters with no effect on any of the variables are not shown. A logarithmic gain of 0 indicates no effect. Strategies involving parameters with red and blue entries are described in the main text.

| Parameters | Dependent Variables |  |  |  |  |
| --- | --- | --- | --- | --- | --- |
| | $M_3$ | $P_8$ | $CM_9$ | $CL_{10}$ | $F$ |
| $\alpha_4$ | 2 | 1 | 1 | 1 | 3 |
| $k_{cat4}$ | 2 | 1 | 1 | 1 | 3 |
| $P_0$ | 2 | 1 | 1 | 1 | 3 |
| $\beta_4$ | -2 | -1 | -1 | -1 | -3 |
| $K_{M4}$ | -2 | -1 | -1 | -1 | -3 |
| $\beta_1$ | 2 | 0 | 0 | 0 | 2 |
| $\beta_5$ | 2 | 1 | 0 | 0 | 2 |
| $K_{M5f}$ | 2 | 1 | 0 | 0 | 2 |
| $U_T$ | 2 | 0 | 0 | 0 | 2 |
| $\alpha_1$ | -2 | 0 | 0 | 0 | -2 |
| $K_2$ | -2 | 0 | 0 | 0 | -2 |
| $\alpha_5$ | -2 | -1 | 0 | 0 | -2 |
| $k_{cat5}$ | -2 | -1 | 0 | 0 | -2 |
| $\alpha_{3max}$ | 1 | 0 | 0 | 0 | 1 |
| $\beta_3$ | -1 | 0 | 0 | 0 | -1 |
| $\alpha_6$ | 0 | 0 | -1 | 0 | 0 |
| $\beta_6$ | 0 | 0 | 1 | 0 | 0 |
| $\alpha_7$ | 0 | 0 | 0 | -1 | 0 |
| $\beta_7$ | 0 | 0 | 0 | 1 | 0 |
| $k_{cat6}$ | 0 | 0 | -1 | 0 | 0 |
| $K_{M6f}$ | 0 | 0 | 1 | 0 | 0 |
| $k_{cat7}$ | 0 | 0 | 0 | -1 | 0 |
| $K_{M7}$ | 0 | 0 | 0 | 1 | 0 |

A number of engineering strategies are contained in Table 2. For instance, interventions increasing the flux through the metabolic pathway without altering the steady state concentration of the potentially toxic metabolic intermediate **P**_**8**_ are shown in color in Table 2 and are graphically represented in Fig. 3A, where each individual arrow represents a different strategy. Note that all these strategies ultimately lead to an increase in the availability of the mRNA molecule **M**_**3**_ and can be categorized into two groups. The first group contains strategies that *directly* increase the synthesis –by either increasing *α*_3max_ or decreasing the binding constant *K*_2_ – and reduce the degradation –by decreasing the rate constant *β*_3_ – of **M**_**3**_. The second group encompasses *indirect* strategies that point at increasing the steady state concentration of the activator form of the transcription factor **U**_**2**_ by modifying rate constants (*α*_1_ or *β*_1_) or by increasing the total pool size of the regulator molecule **U**_**T**_. An analogous analysis can be done to identify strategies increasing the steady state concentration of metabolic intermediates without increasing the pathway flux. We use the **Full System** tab within the **Main Menu** of the user interface of DST3 to demonstrate the validity of these predictions by means of two **titration plots**. In each case, the maximal synthesis rate *α*_3max_ is increased and decreased ten-fold from its nominal operating value of 1 and the effect on the pathway flux **F** (Fig. 3B) and on the steady state concentration **P**_**8**_ (Fig. 3C) is computed for the full system. As predicted by a logarithmic gain of *L* (*F, α*_3max_) = 1 for phenotype **7561**, increasing *α*_3max_ leads to an increase in the steady state flux through the metabolic pathway in the full system. On the other hand, and as indicated by a logarithmic gain of *L* (*P*_8_, *α*_3max_) = 0, increasing or decreasing *α*_3max_ has no effect on the steady state concentration of **P**_**8**_ in the full system. Consequently, *increasing α*_3max_ –which can be experimentally achieved by engineering the promoter region of the polycistronic mRNA or increasing the copy number of the *pca* operon– from its nominal operating value can be used as a strategy to increase the flux through the metabolic pathway without increasing the steady state concentration of **P**_**8**_.

**Figure 3.**
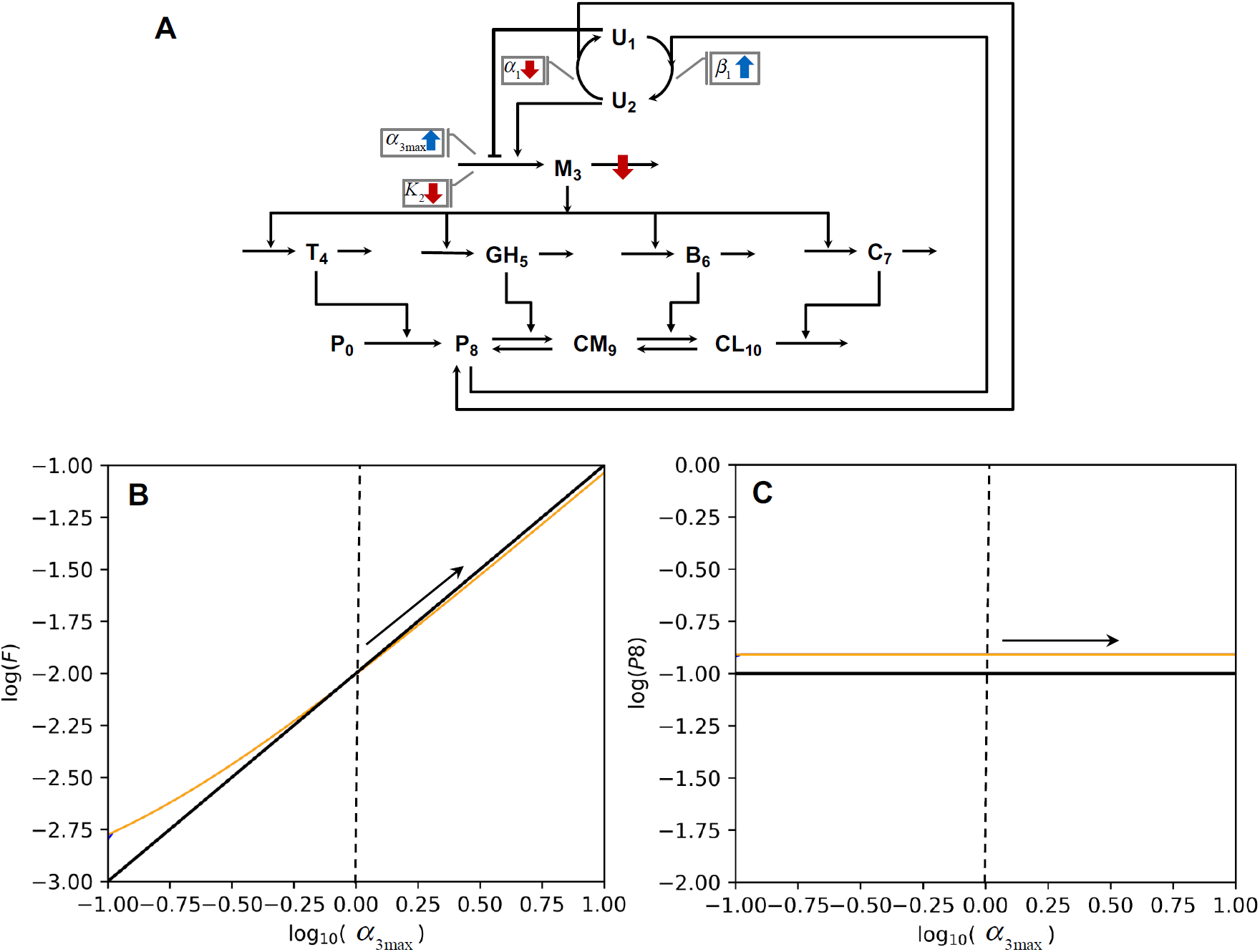
Engineering Strategies for Phenotype 7561. **A)** Five different strategies to increase the flux through the metabolic pathway without increasing the steady state concentration of **P**_**8**_ are shown. Each colored arrow represents an individual strategy. An arrow placed directly over a synthesis or a degradation flux targets its associated rate constant: *α* for synthesis and *β* for degradation. Arrows located within grey boxes usually target kinetic properties of an enzyme or a process. Blue upwards arrows symbolize increase, while red downwards arrows represent decrease. All of the strategies represented in the figure are biologically feasible. However, modifying kinetic properties of a given enzyme or process requires, in most of the cases, a greater experimental effort than modifying its synthesis rate. The effects of perturbing the maximal synthesis rate of the mRNA molecule **M**_**3**_ (*α*_3max_) on the flux through the metabolic pathway **F** (**B**) and the steady state concentration of **P**_**8**_ (**C**) are shown. The operating point of the system is the one depicted in Fig. 2B by the white dot. Vertical dashed lines represent the nominal value of *α*_3max_, from which the system is perturbed. The black solid line represents the behavior of phenotype **7561**. The generation of each **titration plot** for the full system involved numerical integration for 100 different values of *α*_3max_ within the range [0.1 10]. For each point, the system was integrated for 500 time units. To test its stability, the system was integrated for increasing (blue solid line, not shown since covered by orange line) and decreasing *α*_3max_ values (orange line). Since the system follows the same path when integrated forward and backward, it exhibits mono-stable behavior, as predicted by DST3 based on the number of eigenvalues with positive real part for phenotype **7561**. Discrepancies in the location of the solid black and orange line are due to the simplifications made by the Design Space formalism to generate mathematical expressions for phenotype **7561**. However, note that the slope of the black line accurately describes the slope of the orange line.

## 5. Discussion

The Design Space Toolbox v.3.0 offers a variety of advantages over its predecessor version. By distributing the software via a Docker image, the installation process of DST3 is reduced to installing Docker itself. All necessary software dependencies and configurations are already contained within the Docker image, so that users can focus on the actual application of the tools to the analysis of biochemical systems. In line with our goal of improving software usability, DST3 comes with a more stable user interface. By integrating ODE and DAE solvers into the user interface, it is now possible to directly test the accuracy of predictions made by the Design Space formalism from within DST3. Additionally, the capabilities of the computational engine of the Design Space Toolbox were extended. DST3 is now able to analyze biochemical systems containing multiple, nested singularities; something that was out of the reach of previous versions of the toolbox. We demonstrated the utility of DST3 by analyzing an integrated biochemical system consisting of a signaling cascade, a gene circuit and a metabolic pathway. The system’s topology encoded a cycle, a conservation relationship and the potential to exhibit blow-up behavior.

We applied a recently developed phenotype-centric modeling strategy (Valderrama-Gómez et al 2018, Lomnitz and Savageau 2015) to identify a stable and globally robust operating point of the system. This process involved listing the phenotypic repertoire and filtering it for phenotypes of interest. From a total of 3722 possible phenotypes, 384 were found to be non-pathological, 192 of these physiological phenotypes were responsive to changes in the concentration of environmental substrate protocatechuate (**P**_**0**_), 64 of the responder phenotypes were found to be inducible. The latter fell into three groups based on the steepness of the induction characteristic: L(M_3_,P_0_)=2 with 32 phenotypes, L(M_3_,P_0_)=4 with 28, and L(M_3_,P_0_)=6 with 4. When compared on the bases of three criteria, global robustness, energy efficiency and toxicity, the first group was best and the third group was worst. An analysis of the volume of the 64 phenotypes with desired properties revealed that their combined volume only accounted for 5.61×10^−5^ % of the total volume of all non-pathological phenotypes identified by DST3 (see Table 1). When pathological phenotypes were considered (phenotypes exhibiting a blowing behavior), this value decreased to 4.50×10^−24^ %. This suggests that desirable phenotypes will have to be actively selected for by nature, since the vast majority of parameter values chosen at random (increased entropy) would produce few desirable phenotypes.

These figures highlight the power of DST3 and the phenotype-centric modeling strategy it enables. Finding the reported operating point for the representative phenotype **7561** and characterizing its robustness and associated boundaries in a 30-dimensional parameter space by means of parameter sampling would have been computationally expensive and impractical. Indeed, current methods based on the ensemble modeling approach (Tran et al 2008) for robustness analysis (Lee et al 2014) involve computationally expensive dense parameter sampling and numerical integration by ODE solvers for stability assessment. These approaches require a long computational time for large model ensembles, and they do not allow for a rigorous identification of stability boundaries. On the other hand, the Design Space formalism decomposes the parameter space into a set of polytopes, biochemical phenotypes, whose boundaries and properties are well defined. DST3 not only identifies these phenotypes, but it also allows the automatic prediction of nominal parameter sets for their realization. This greatly facilitates deterministic simulations of the full system that require parameter values, as demonstrated in panels C and E of Fig. 2 and panels B and C of Fig. 3. Similarly, stochastic simulations, which also require parameter values, can benefit from the innovations offered by DST3.

DST3 predictions regarding steady states, stability and blowing behavior were accurate, as demonstrated by time course, titration and trajectory plots generated for the full system. By finding strategies to increase the flux through the metabolic pathway of the system without increasing the steady state concentration of an intermediate metabolite, we aimed at showing a glimpse of the potential that the Design Space formalism has to offer to the field of *rational* Metabolic Engineering (Bailey 1991). Further potential applications relate to the ability of DST3 to correctly identify and characterize blowing phenotypes, which are commonly found in metabolic systems. Often, in the process of strain development, intermediate strains are generated, in which a given intermediate metabolite excessively accumulates or is totally consumed, thus generating a metabolic imbalance within the cell. This decreases strain fitness and can ultimately lead to cellular death (Dahl et al. 2013, George et al 2014, Alonso-Gutierrez et al. 2017). DST3 is able to identify regions in the parameter space leading to metabolic imbalances and to provide clues to rectify these phenotypes. For instance, consider the operating point of the system shown in Fig. 2D, which is located within the blowing phenotype 7718.1. Inspection of the Design Space plot around this phenotype indicates that increasing the value of k_cat5_ to values larger than 0.1 would place the operating point of the system within phenotype 7705, 7561, 649 or 1945_2, all of which exhibit a stable, non-pathological steady state. The specific location of the operating point within any of these phenotypes will depend on the extend of the increase of the parameter k_cat5_. Alternative strategies to rectify the pathological behavior of an operating point located within phenotype 7718.1 include increasing the amount of the PcaGH enzyme by cloning its gene sequence on a controllable plasmid or engineering its ribosomal binding site.

The application of mechanistic models for the identification of metabolic engineering strategies has been rather limited. This has been mainly caused by a lack of knowledge of associated parameter values. As a consequence, constraint-based modeling has been the method of choice applied to rationally guide metabolic engineering strategies (Valderrama-Gómez 2017). By enabling a parameter-free, mechanistic, phenotype-centric modeling strategy, the Design Space formalism and associated toolbox offers enormous potential for the field of metabolic engineering.

Elucidating biological design principles is another important area for application of the Design Space formalism that was not explored in this work due to space limitations. In the Design Space, boundaries delimiting biochemical phenotypes are linear functions of the system’s parameters in logarithmic coordinates. Thus, design principles can be readily identified in the form of mathematical inequalities involving the parameters of the system. These ideas were applied by Fasani and Savageau (2013) to study properties of toxin-antitoxin systems, which have been linked with the medically relevant persister phenotype exhibited by certain bacterial strains. The study revealed factors affecting the frequency of persisters in the population, such as the overall number of toxin-antitoxin modules and the size and position of the bistable region, a property emerging from the system’s architecture.

There are many examples of systems that appear to perform the same function, and yet they exhibit radically different genomic architectures, the reasons for which are poorly understood. An example is provided by the protocatechuate degradation pathway studied in this work. It is one of the two branches of the β-ketoadipate pathway, a chromosomally encoded convergent pathway for aromatic compound degradation that is widely distributed in soil bacteria and fungi. Enzyme studies suggests that the pathway is highly conserved in diverse bacteria; however, its regulation and gene organization differ greatly (Harwood and Parales 1996). For instance, the pathway genes from *Pseudomonas aeruginosa* and *P. synrigae* are arranged in three and four different clusters, respectively. By contrast, all genes are arranged in a single cluster in *Acinetobacter* sp. ADP1 (studied in this work) and in *P. fluorescens* (Jiménez et al. 2002). It has been suggested that evolutionary processes have shaped moldable aspects of the β-ketoadipate pathway to optimally serve diverse lifestyles of bacteria (Harwood and Parales 1996). DST3 could be used to compare and contrast inherent aspects of each system, such as its dynamic properties, induction characteristics and tradeoffs regarding energy and toxicity, thus potentially allowing the elucidation of underlying design principles used by nature to create the alternative genomic architectures observed in organisms with different environments and lifestyles.

## 6. Materials and Methods

### Method Details

Here, we show how various computational tools are integrated to create DST3. We go on to explain how Docker images and containers can be used to access DST3 on virtually any operating system. Then, we briefly describe two components of DST3 that can be used to access the computational capabilities of DST3: its user interface and its Python module. Refer to Ipython-notebooks contained in the Docker image under **/Tutorials/Tutorial_DST3** for a detailed description of the user interface.

### 6.1 Design Space Toolbox v.3.0

Innovations contained in DST3 aim at improving three key aspects of the software: utility, usability and portability. By further developing the C library of DST3 to allow for the automatic identification and mathematical characterization of various types of singularities, we increased the scope of systems that can be analyzed by DST3, thus improving its *utility*. By enhancing stability and functionality of the IPython-based user interface of DST3, we increased software *usability* for users with limited programming knowledge. For advanced users, we generated a python module that runs on python 3.7.3 and can be integrated into customized programs. Due to its various external software dependencies, DST2 suffered from a limited portability. We addressed this issue by packaging DST3 into a Docker image. This effectively renders Docker the only software dependency necessary to run DST3 and guarantees *portability* across major operating systems.

#### 6.1.1 Technology overview

Three main components make up DST3: A C library, a Python package and a user interface (Fig. S1). All three components are interconnected, with the C library being the computational engine that performs most of the numerical analyses. The Python package was designed to provide high-level access to the C library, making further software development simpler and faster. The user interface was built using IPython widgets and accesses the C library through the Python module.

Four steps are required to install and access all components of DST3:

1. Install Docker on your operating system
2. Download the latest DST3 image by typing the following command on a Terminal or Prompt Window:

~~~
*docker pull savageau/dst3*
~~~ This will download the latest stable version of DST3 running on Python 2.7.3, for which a user interface is available. For advanced users, a Docker image of DST3 running on Python 3.7.3 (only the Python module is available) can be downloaded instead by typing:

~~~
*docker pull savageau/dst3:python3*
~~~
3. Start a Docker container to access DST3 by typing the following command on a Terminal or Prompt window:

~~~
*docker run -d -p 8888:8888 savageau/dst3*
~~~ That command will create a container without access to the files of the host computer. Files created within the container will be lost after the container is stopped. In order to grant access to files on the host computer, the previous command should be complemented with the flag *--mount*:

~~~
*docker run -d -p 8888:8888 --mount
type=bind,source=/Users,target=/Documents/host savageau/dst3*
~~~ Windows users should use:

~~~
*docker run -d -p 8888:8888 --mount
type=bind,source=//c/Users,target=/Documents/host
savageau/dst3*
~~~
4. Access DST3 by opening the following address on any internet browser: http://localhost:8888/

#### 6.1.2 DST3 User Interface

DST3 comes with an updated and more stable Ipython-based user interface. Fig. S2 presents a hierarchical overview of the different menus available. The gray box represents the initial menu, from which the functionality of DST3 can be accessed. The **About** menu provides general information about the software, including its version, developers and an option to report bugs. Syntax rules for equations are also contained in this menu. Analyses supported by DST3 can be accessed from the **Main Menu**. Results, in form of figures and tables, are managed by the **Figures** and **Tables** menu, respectively. The **System** tab contains general information about the specific set of differential equations subject to analysis. It includes the name of the system, the total number of cases and its system signature.

Sub-menus and tabs contained in **Main Menu** include the following:

##### Phenotypic Repertoire

This window allows the user to list and filter biochemical phenotypes – cases – according to their validity, case signature, log-gain values, number of eigenvalues with positive real part and volume. The resulting phenotypic repertoire can be exported to a `.xlsx` file or saved into the **Tables** menu if desired.

##### Analyze Case

This tab creates the full analysis of a case referenced by its case number or case signature. This analysis includes the S-system equations that mathematically define the case, conditions that need to be fulfilled for its validity, its steady state solution – if it exists – and boundary constraints. For cases with a steady state solution, logarithmic gains for dependent variables with respect to independent variables and parameters are calculated and reported. For valid cases, a bounding box for the corresponding high-dimensional polytope is also provided. Additionally, it is possible to estimate a set of parameter values located within this high-dimensional polytope through linear programming. Global tolerances can be calculated from that nominal parameter set or from any other parameter set within the polytope by calculating respective lower and upper bounds for each parameter. The ***Analyze Case*** window also provides eigenvalues for the respective S-system. Note that individual tables generated by the ***Analyze Case*** tab can be saved into the **Tables** menu.

##### Case Intersections

For a given set of cases, the ***Case Intersections*** window indicates if there is a region in parameter space where the cases overlap. If this region exist, global tolerances are reported for a parameter set located within the intersecting polytope.

##### Co-localizations

For a given set of cases and so-called slice variables, the ***Co-localizations*** tab indicates if regions of validity for each case exist within the given slice. If the co-localization is valid, global tolerances are reported. Additionally, design space plots can be generated to visualize the co-localization of the cases.

##### Create Plot

This window allows the generation of multiple plots, which are useful in characterizing the design space and its predictions for stability, steady state concentrations, fluxes, etc. Even though an explorative characterization of the design space is possible via the *interactive* design space plot, functionalities contained in the ***Create Plot*** window are more effective when used to characterize a specific region of interest in the parameter space. This region can be usually found by combining functionalities of the ***Phenotypic Repertoire*** and ***Analyze Case*** windows. Plots generated by this window can be saved in the menu **Figures.**

##### Full System

It allows the characterization of the temporal response (Time Courses plot) of the original set of differential equations under study, as well as the analysis of its stability properties (Titration and Trajectories Plots). This characterization is useful to assess the accuracy of predictions made using the S-system approximations. Analysis of systems of differential algebraic equations (DAE) is performed through the python package Assimulo (Andersson et al 2015). Plots generated by this window can be saved in the menu **Figures.**

#### 6.1.3 DST3 Python Module

The capability of the DST3 C library to identify special phenotypes can be enabled by changing default values of arguments passed to classes `Equations` and `DesignSpace`. Both classes are contained in the DST3 Python Package `dspace` and play a central role in generating Python objects involved in any computational Design Space analysis, as shown in Fig. S3. The input to the class `Equations` consists of a machine-readable string representation of the system of equations describing the dynamics of the network. A string representation that can be parsed by the computational engine of DST3 should be compliant with following syntax rules:

1. Each equation has to be explicitly stated as:
  a. A differential equation, where the “.” operator denotes the derivative with respect to time.
  b. An algebraic constraint, where the left-hand side is either a variable or a mathematical expression. Auxiliary variables associated with the constraint must be explicitly defined (unless the left-hand side is the auxiliary variable). Algebraic constraints can be used to represent conservation constraints. In that case, they should be placed after regular algebraic constraints. Additionally, they should follow the form dictated by Eq. S21. When using the Python module, associated auxiliary variables should be explicitly declared as ‘Xci’, with *i* = 1,…, *n*_*cr*_. This definition is not necessary when using the DST3 user interface. In any case, variable names ‘Xci’, with *i* = 1,…, *n*_*cr*_ are reserved for the computational engine and should not be contained in system of equations defined by the user.
2. Multiplication is represented by the “*” operator.
3. Powers are represented by the “^” operator.
4. Architectural constraints are defined as inequalities, where both sides of the inequality are products of power-laws.

In order to exemplify the generation of valid machine-readable string representations and the usage of the DST3 Python module, we calculate valid cases for three different synthetic biochemical systems, each one exhibiting a different type of singularity. For each system, five lines of Python code are presented and discussed. Computational steps involve in each case:

1. Importing the DST3 python module `dspace`,
2. Defining a string representation for the system,
3. Generating an `Equations` object,
4. Generating a `DesignSpace` object,
5. Generating a list of valid cases.

##### Cycles

The synthetic network under analysis is described by Eqs. S1-S3. Generating a string representation of this system is straightforward and results in a list of three strings, one for each differential equation, as shown in `line 2` of the snippet below.

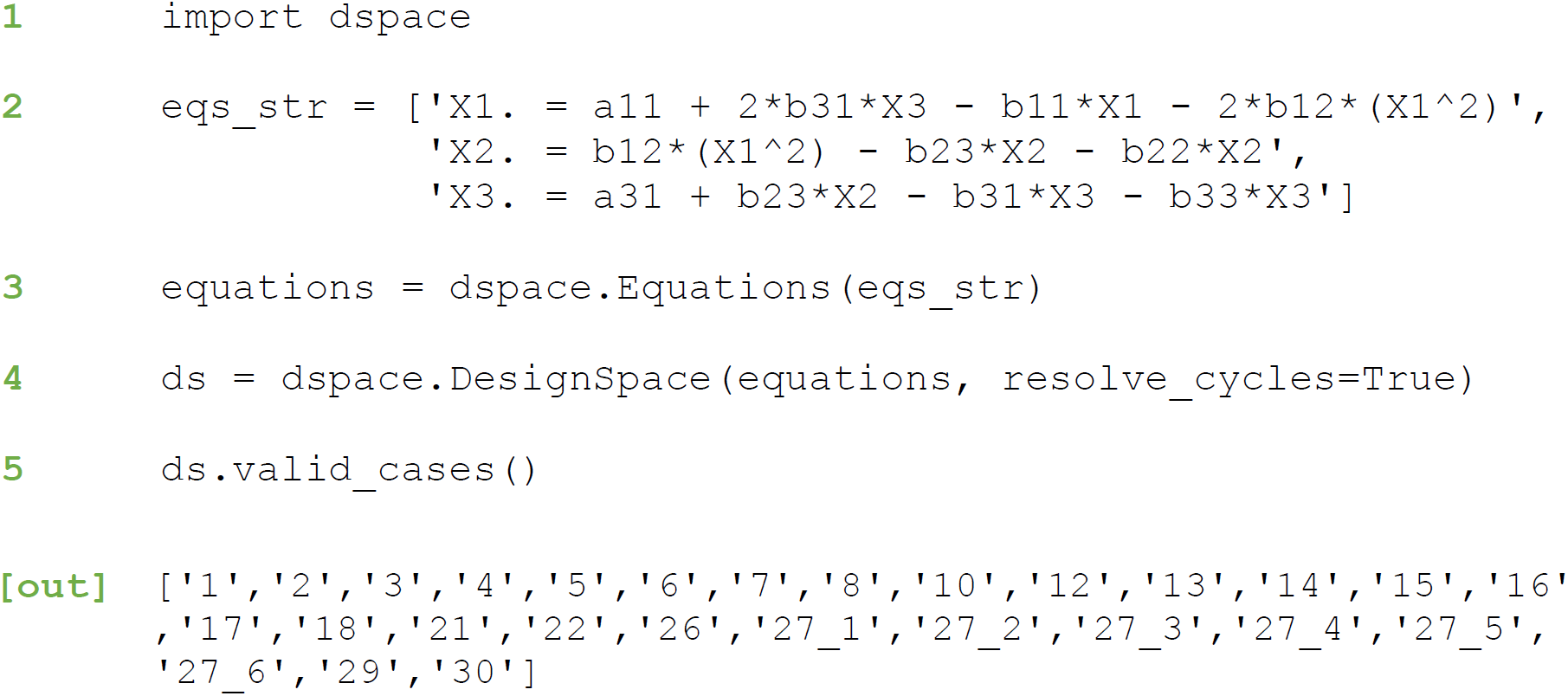

Since the system does not contain any auxiliary variable, the object `eq_string` is passed as sole positional argument to the class `dspace.Equations()` to generate the `Equations` object, which is stored in the variable `equations`. In order to identify and resolve the cycle encoded within this system, the key argument `resolve_cycles` is set to `True` and passed along with the `equations` object to the class `dspace.DesignSpace()` to generate a `DesignSpace` object which is stored in the variable `ds`. A list of valid cases is generated through the method `valid_cases()` of the `ds` object. Note that cases `27_1, 27_2,…, 27_6` result from resolving the cyclical case 27 (refer to Table S1).

##### Conservations

As discussed before, the system described by Eqs. S22-S25. contains a conservation constraint among its constituent pools, as defined by Eq. S25. According to the DST3 syntax rules, this conservation relationship can be explicitly defined as an algebraic constraint and should be placed in the last position of the string representation of the system (see `line 2` of the snippet below). Since the Python module is being used to analyze this system, an associated auxiliary variable needs to be explicitly defined, as shown in `line 3`.

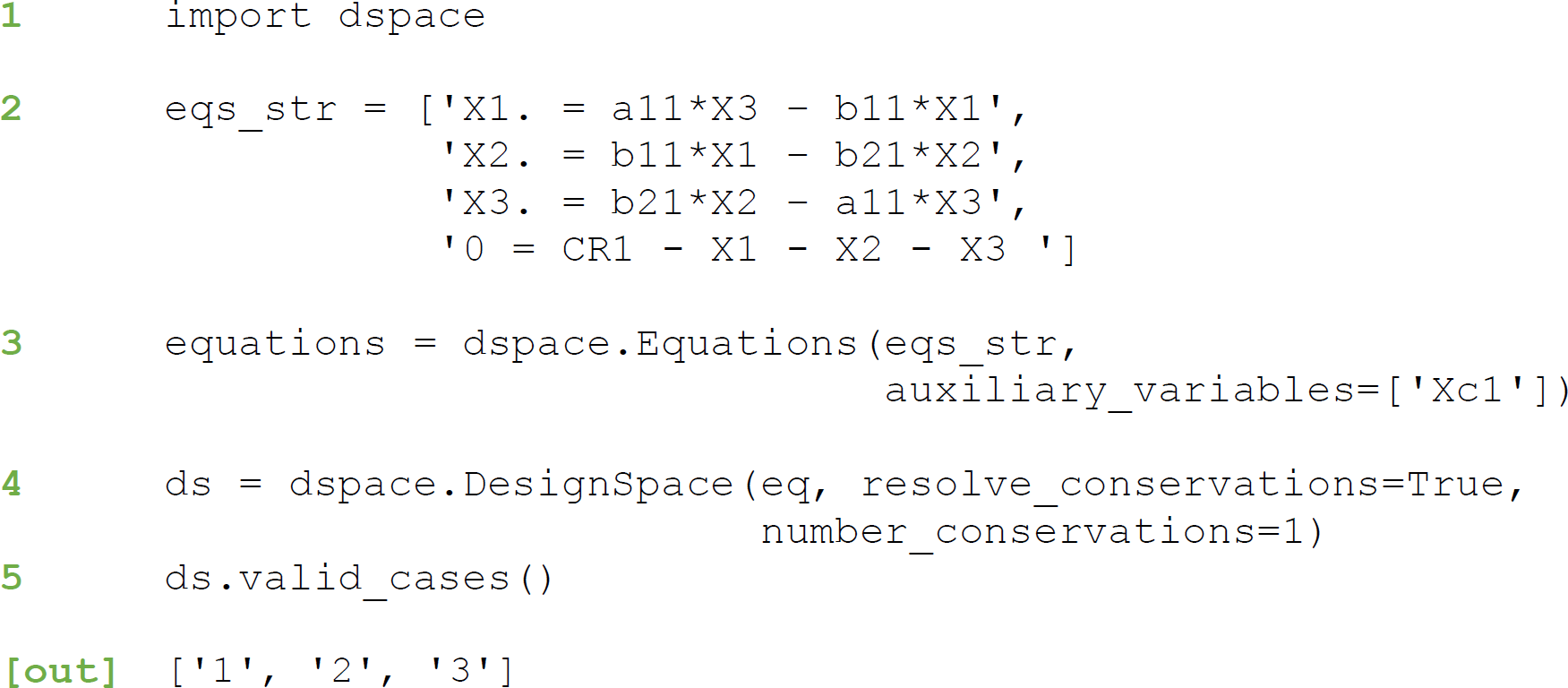

The computational engine of DST3 is informed about the conservation constraint by using two key arguments: `number_conservations=1` and `resolve_conservations=True.` Valid cases can be printed by means of the method `valid_cases()` of the `ds` object.

##### Metabolic Imbalances

The last example consists of a system containing one differential equation and one algebraic constraint, as defined by Eqs. S26-S27. For certain parameter values, this system has the potential to exhibit a blow-up for variable *X*_1_. The class `dspace.Equations` is informed about the algebraic constraint contained in the string representation by using the key argument `auxiliary_variables=[‘D’].` Similarly, the DST3 computational engine is configured to check for blow-ups by using the key argument `resolve_instability=True`, as shown in the snippet below:

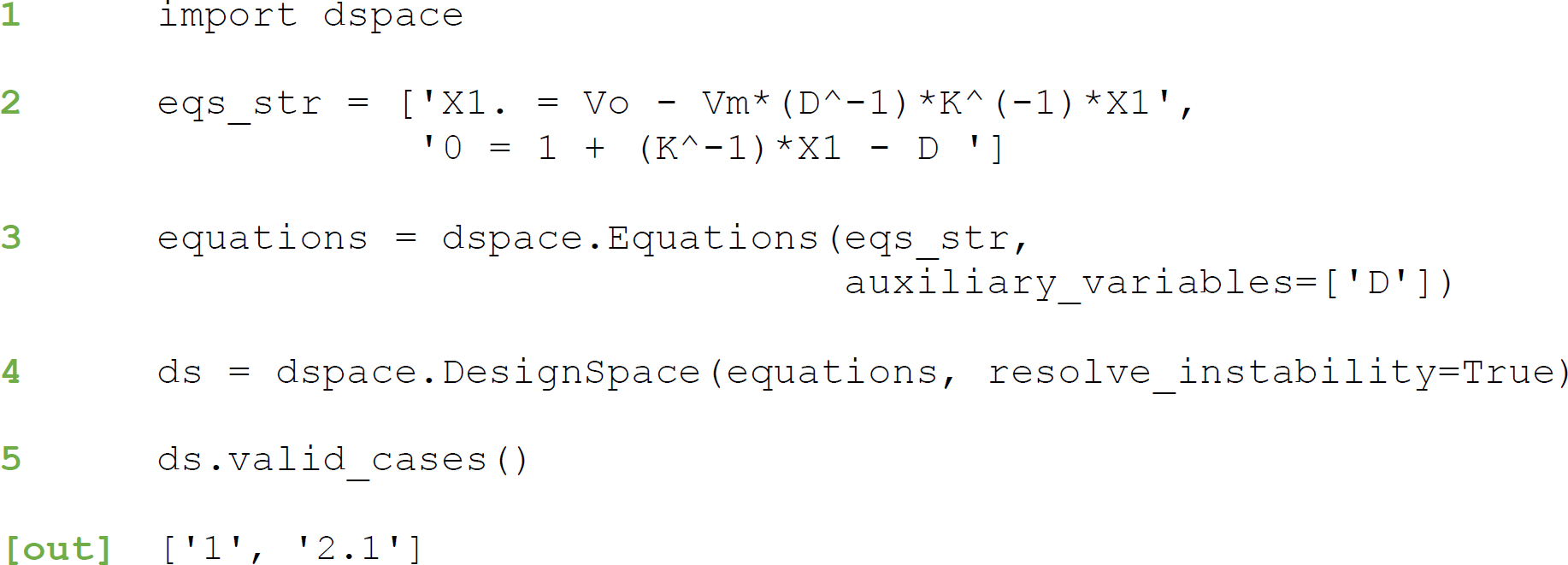

Valid cases can be customarily printed using the method valid_cases()of the ds object. Note that the case identifier 2.1 refers to the sub-case 2.1 of Table S3.

### 6.2 Data and Code Availability

The Docker images used by DST3 are freely available at https://hub.docker.com/r/savageau/dst3

### 6.2 Additional Resources

A tutorial is available as various IPython notebooks within the DST3 docker image under **/Tutorials/Tutorial_DST3**

## 7. Acknowledgements

This work was supported in part by a grant from the US National Science Foundation Grant number MCB 1716833.

## 8. Author contributions

Conceptualization, M.A.S., R.A.F., J.G.L. and M.A.V.; Methodology, M.A.S. and M.A.V.; Software, J.G.L and M.A.V; Validation, M.A.V. and M.A.S.; Investigation, M.A.S and M.A.V.; Writing, M.A.V. and M.A.V.; Supervision, M.A.S., Funding Acquisition, M.A.S.

