## Supplemental Information for "Mechanistic Modeling of Biochemical Systems Without A Priori Parameter Values Using the Design Space Toolbox v.3.0"

### 1. Strategies for Treating Three Types of Singularities

#### *1.1 Cycles are resolved by considering global dominance equations.*

We start by setting up Eqs. S1 to S3 to describe the change in time for the concentration of each chemical species shown in Fig 1A in the main text. Mass action kinetics are used to generate rate laws describing the flux through each reaction of the network. The resulting expressions are then combined by means of Kirchhoff's node law to generate balance equations for each metabolite in the network.

$$\frac{dX_1}{dt} = \alpha_{11} + 2\beta_{31}X_3 - \beta_{11}X_1 - 2\beta_{12}X_1^2 \tag{S1}$$

$$\frac{dX_2}{dt} = \beta_{12}X_1^2 - \beta_{23}X_2 - \beta_{22}X_2 \tag{S2}$$

$$\frac{dX_3}{dt} = \alpha_{31} + \beta_{23}X_2 - \beta_{31}X_3 - \beta_{33}X_3. \tag{S3}$$

The Design Space formalism can be applied to decompose this set of equations into different cases, each having a unique set of dominant terms and being valid within a specific region in parameter space. Eqs. S4 – S6 represent one of those cases, which is defined by the case signature [22 11 21] and the case number 27.

$$\frac{dX_1}{dt} = 2\beta_{31}X_3 - 2\beta_{12}X_1^2 \tag{S4}$$

$$\frac{dX_2}{dt} = \beta_{12}X_1^2 - \beta_{23}X_2 \tag{S5}$$

$$\frac{dX_3}{dt} = \beta_{23} X_2 - \beta_{31} X_3 . \qquad \text{(S6)}$$

Necessary conditions for these terms to be dominant are described by Eqs. S7 – S11:

$$2\alpha_{11}^{-1}\beta_{31}X_3 > 1 \qquad \text{(S7)}$$

$$2\beta_{11}^{-1}\beta_{12}X_1 > 1 \qquad \text{(S8)}$$

$$\beta_{23}\beta_{22}^{-1} > 1 \qquad \text{(S9)}$$

$$\beta_{23}\alpha_{31}^{-1}X_2 > 1 \qquad \text{(S10)}$$

$$\beta_{31}\beta_{33}^{-1} > 1 \ . \qquad \text{(S11)}$$

By setting the left-hand side of Eqs S4-S6 to zero, taking logarithms and rearranging, one obtains Eq. S12, which exhibits the form of Eq. 8 in the main text:

$$\begin{pmatrix} -2 & 0 & 1 \\ 2 & -1 & 0 \\ 0 & 1 & -1 \end{pmatrix} y_D + \begin{pmatrix} -1 & 1 & 0 \\ 1 & 0 & -1 \\ 0 & -1 & 1 \end{pmatrix} y_I = \begin{pmatrix} 0 \\ 0 \\ 0 \end{pmatrix} , \qquad \text{(S12)}$$

with $y_D^T = \begin{pmatrix} \log X_1 & \log X_2 & \log X_3 \end{pmatrix}$ and $y_I^T = \begin{pmatrix} \log\beta_{12} & \log\beta_{31} & \log\beta_{23} \end{pmatrix}$. Visual inspection of the matrix $A_D$ reveals the presence of a linear dependency among its rows: $row_1(A_D) = -row_2(A_D) - row_3(A_D)$. This causes matrix $A_D$ to be rank deficient [$rank(A_D) = 2$] and prevents the computation of a unique solution for $y_D$. Since $rank(A_D) = rank(A_I)$, the system of algebraic equations is consistent and a solution (or set of solutions) can be found by applying the design space formalism to a sub-system defined by an extended set of equations. The modification of the system consists in adding a so-called *global dominance* equation, which describes a mass balance around each cycle present in the system. The following steps are involved in generating the extended set of equations:

1. Identify set(s) of *cyclical variables* $C$ by computing the null space of matrix $A_D$. Cyclical variables are characterized by non-zero row entries in the null space matrix. Note that the number of cycles contained in $A_D$ is defined by the number of columns of the null space: $col(Null(A_D))$. For the specific case being analyzed one obtains: $\left[Null(A_D)\right]^T = \left[\begin{array}{ccc} 1 & 1 & 1 \end{array}\right]$, meaning that the system contains one single cycle, with the set of cyclical variables $C = \left\{\begin{array}{ccc} X_1, & X_2, & X_3 \end{array}\right\}$.
2. Set up a *global dominance* equation for each cycle. This equation is a mathematical representation of a mass balance around a given cycle in steady state. In the case of the network considered in Fig 1A, the global dominance equation will be a function of fluxes entering and leaving the control volume delimited by the blue rectangle in Fig 1B, i.e, fluxes governed by rate constants $\alpha_{11}, \alpha_{31}, \beta_{11}, \beta_{22}$ and $\alpha_{33}$. Global dominance equations are constructed by weighting those fluxes using coefficients obtained from the null space of matrix $A_D^T$. In this specific case one obtains: $\left[Null(A_D^T)\right]^T = [\ 1 \quad 2 \quad 2\ ]$, which yields the following global dominance equation: $0 = \alpha_{11} - \beta_{11}X_1 - 2X_2\beta_{22} + 2\alpha_{31} - 2X_3\beta_{33}$.
3. Generate extended sub-system by introducing the global dominance equation(s). Eqs. S13-S16 along with the conditions defined by Eqs. S7-S11 define the extended sub-system that should be used to resolve the cyclical case generated by Eqs. S4-S6.

$$\frac{dX_1}{dt} = 2\beta_{31}X_3 - 2\beta_{12}X_1^2 \tag{S13}$$

$$\frac{dX_2}{dt} = \beta_{12}X_1^2 - \beta_{23}X_2 \tag{S14}$$

$$\frac{dX_3}{dt} = \beta_{23}X_2 - \beta_{31}X_3 \tag{S15}$$

$$0 = \alpha_{11} + 2\alpha_{31} - \beta_{11}X_1 - 2\beta_{22}X_2 - 2\beta_{33}X_3 . \tag{S16}$$

DST3 implements steps 1 to 3 in a recursive fashion to resolve multiple and nested cycles. Once the extended sub-system has been set up, the design space formalism can be applied to identify valid sub-cases resolving the cyclical case. Table S1 shows S-system equations for each one of the six valid cases generated from the extended sub-system. Note the special form of these equations. S-systems originating from a dominance analysis on the global dominance equation (Eq. S16) are used to replace the differential equation for the pool with the dominant efflux. For instance, the expression $\alpha_{11} - \beta_{11} X_1$, which is obtained when the first positive and first negative term in Eq. S16 are dominant, is used to replace the differential equation for $X_1$ (refer to sub-case 27_1 in Table S1). Additionally, this expression is scaled to match the coefficient of the negative term in the full system (i.e., the original set of equations). Consider for instance the expression $\alpha_{11} - 2\beta_{33} X_3$, which is obtained when the first positive and third negative term of Eq. S16 are dominant. Since the original coefficient of the negative term is 1 (see Eq. S3), the scaled expression $\frac{1}{2}\alpha_{11} - \beta_{33} X_3$ is used to construct sub-case 27_3 of Table S1.

*1.2 Conserved moieties are handled by considering the total size of conserved pools.*

Consider the simple system in Fig. 1C in the main text with three components linked by a conservation relationship. Mass balance equations can be set up for each metabolite using the rate laws shown in Fig. 1C:

$$\frac{dX_1}{dt} = r_1 - r_2 \tag{S17}$$

$$\frac{dX_2}{dt} = r_2 - r_3 \tag{S18}$$

$$\frac{dX_3}{dt} = r_3 - r_1 . \tag{S19}$$

Balance Eqs. S17-S19 can be compactly expressed in matrix form to yield Eq. S20:

$$\frac{dX}{dt} = Sr , \tag{S20}$$

with $S$ being the stoichiometric matrix and $r$ a vector of rate laws describing the flux through each reaction. The number of conservations $n_{cr}$ within the system is given by: $n_{cr} = col(Null(S^T))$, which equals 1 for the system under consideration.

Conservation relationships can be mathematically described as linear dependencies among metabolite pools. They can be expressed in matrix form as:

$$0 = CR - Null(S^T)^T \times X. \qquad \text{(S21)}$$

$CR$ is a vector of independent variables and $CR_i$ expresses the total pool size of each conservation. $X$ represents a vector of concentration pools. Applying Eq. S21 to the system defined by Eqs. S17-S19 yields: $0 = CR_1 - X_1 - X_2 - X_3$. DST3 is able to handle biochemical systems with single or multiple conservation relationships. For the system depicted in Fig. 1C, Eqs. S22-S25 represents an appropriate set of equations that can be analyzed by DST3 given the conservation relationship provided:

$$\frac{dX_1}{dt} = \alpha_{11} X_3 - \beta_{11} X_1 \qquad \text{(S22)}$$

$$\frac{dX_2}{dt} = \beta_{11} X_1 - \beta_{21} X_2 \qquad \text{(S23)}$$

$$\frac{dX_3}{dt} = \beta_{21} X_2 - \alpha_{11} X_3 \qquad \text{(S24)}$$

$$0 = CR_1 - X_1 - X_2 - X_3 \qquad \text{(S25)}$$

The analysis of this system using the Design Space formalism involves the usual generation of cases by picking dominant terms for each of the equations. To handle the singularity generated by the conservation encoded in the system, DST3 eliminates the differential equation(s) corresponding to the dominant negative term of each conservation relationship. This generates cases with S-systems that are deficient in $n_{cr}$ differential equations. Table S2 contains three cases that result from applying the design space formalism to the system defined by Eqs S22-S25. Note that each case is defined by only two differential equations and one algebraic constraint. In order to capture this special way of constructing case equations, the indices of the differential equation being deleted are set to zero in the case signature. Case 3 for instance, in which the differential equation for pool $X_3$ is missing, has a case signature of [11 11 **00** 13] to reflect this fact.

Note the similarity between the system topology of the cyclical case 27 (Fig. 1B) and the conserved system of this section (Fig. 1C). In both instances, metabolites $X_1$, $X_2$ and $X_3$

interact in a cyclical fashion to introduce a linear dependency among their pools that renders their $A_D$ matrix singular. For cyclical cases (e.g. case 27), this dependency is eliminated by means of a global dominance equation, which represents a mass balance around the cycle. This strategy cannot be applied for conserved moieties, because they are not synthesized, degraded or exchanged with the environment (Haraldsdóttir and Fleming, 2016). From a mass balance perspective, this implies that fluxes entering or leaving the conservation do not exist, as exemplified in Fig. 1D. The singularity is thus eliminated by replacing the differential equation for one of the metabolites involved in the conservation by an algebraic constraint that contains an additional independent parameter ( $CR_i$ ).

*1.3 Metabolic Imbalances are treated by considering knife-edge conditions*

Given the system shown in Fig. 1E and described by Eqs. 11-12 in the main text, consider the equations for case 2 with signature [11 21]:

$$\frac{dX_1}{dt} = v_0 - V_M D^{-1} K_M^{-1} X_1 \quad \text{(S26)}$$

$$0 = K_M^{-1} X_1 - D \ , \quad \text{(S27)}$$

and its associated dominance condition:

$$K_M^{-1} X_1 > 1 . \quad \text{(S28)}$$

Substituting the algebraic constraint (Eq. S27) into the differential equation for $X_1$ (Eq. S26) yields the following dynamical system:

$$\frac{dX_1}{dt} = v_0 - V_M . \quad \text{(S29)}$$

Setting the left-hand side of this equation to zero, taking logarithms of both sides and rearranging in matrix notation analogously to Eq. 8 in the main text results in:

$$\begin{bmatrix} 0 \end{bmatrix} y_D + \begin{bmatrix} 1 & -1 \end{bmatrix} y_I = \begin{bmatrix} 0 \end{bmatrix}, \tag{S30}$$

with $y_D = \begin{bmatrix} \log X_1 \end{bmatrix}$ and $y_I^T = \begin{bmatrix} \log v_0 & \log V_M \end{bmatrix}$. Since $rank(A_D) < rank(A_I)$, the system does not have a steady state solution. Indeed, Eq. S29 only provides a consistency condition for the concentration $X_1$ to remain unchanged over time: $0 = v_0 - V_M$. We will refer to this kind of constraint as a *knife-edge condition*. In general, we are interested in the behavior of the system when knife-edge conditions are not satisfied, i.e., $v_0 \neq V_M$. Violating the knife-edge condition in a specific direction implies an extreme value for $X_1 : X_1 \to \infty$ or $X_1 \to 0$. The validity of either situation is assessed by checking the validity of the associated dominance conditions, as shown in Table S3. Taken together, these results indicate that for the system shown in Fig. 1E, the concentration of the pool $X_1$ will steadily increase over time, i.e., it will *blow up* if independent variables fulfill the conditions $K_M^{-1} X_1 > 1$ and $v_0 V_M^{-1} > 1$.

To introduce further concepts required to establish a general framework for the treatment of cases for which $rank(A_D) < rank(A_I)$, let us now consider the system described by Eqs. S31-S32:

$$\frac{dX_1}{dt} = \alpha_1 - \alpha_2 \tag{S31}$$

$$\frac{dX_2}{dt} = \alpha_2 - X_1, \tag{S32}$$

which when rearranged and expressed in matrix notation to resemble the form of Eq. 8 in the main text yields:

$$\begin{pmatrix} 0 & 0 \\ -1 & 0 \end{pmatrix} y_D + \begin{pmatrix} 1 & -1 \\ 0 & 1 \end{pmatrix} y_I = \begin{pmatrix} 0 \\ 0 \end{pmatrix}, \tag{S33}$$

with $y_D^T = \begin{bmatrix} \log X_1 & \log X_2 \end{bmatrix}$ and $y_I^T = \begin{bmatrix} \log \alpha_1 & \log \alpha_2 \end{bmatrix}$. Since $rank(A_D) = 1$ and $rank(A_I) = 2$, the system is not consistent and does not have a valid steady state solution. The structure of the matrix $A_D$ indicates the existence of two knife-edge conditions, which give rise to four sub-cases, as shown in Table S4. In general, the number of subcases to be tested equals $2^{n_{knife}}$, with $n_{knife}$ the number of knife edges present in the system. Out of four possible sub-cases, only sub-case 2 and sub-case 3 are valid because of the way in which they violate both knife-edge conditions is consistent. The fact that the matrix $A_D$ has one degree of freedom ( $row(A_D) - rank(A_D) = 2 - 1 = 1$ ), implies a relationship between the two knife edge conditions. Indeed, violating the knife-edge condition $\alpha_1 = \alpha_2$ in either direction sets an extreme value for $X_1$ . This in turn dictates the way in which the second knife edge $\alpha_2 = X_1$ will be violated, thus setting an extreme value for $X_2$. Since the structure of the matrix $A_D$ prevents $X_2$ from being calculated via matrix operations after an extreme value for $X_1$ has been set, we opt for testing the validity of each possible sub-system via linear programming.

More generally, three steps are involved in resolving cases for which $rank(A_D) < rank(A_I)$:

1. Merge auxiliary variables into differential equations to obtain dynamical systems without algebraic constraints.
2. Identify the number and identity of knife-edge conditions as a function of the degrees of freedom ( $n_{freedom}$ ) of matrix $A_D$ and the number of zeros in its diagonal ( $n_{zeros}$ ):
    a. If $n_{freedom}$ is equal to $n_{zeros}$, then $n_{knife} = n_{freedom}$ and knife edge conditions correspond to balance equations for pools with a zero entry in the diagonal of $A_D$
    b. If $n_{freedom} < n_{zeros}$, then $n_{knife} = n_{zeros}$ and knife edge conditions correspond to balance equations for pools with a zero entry in the diagonal $A_D$
    c. If $n_{freedom} > n_{zeros}$, then $n_{knife} = n_{freedom}$ and knife edge conditions correspond to balance equations for $n_{knife}$ randomly selected pools.

3. Construct and test the validity of $2^{n_{knife}}$ different linear programs, each violating knife edge conditions in a unique way. Note that the test for validity should include dominance conditions associated with the case under analysis.

### 2. **Analysis of a Biochemical System Exhibiting Multiple Singularities**

#### *2.1 Differential Equations for the Biochemical System*

*Transcription factor:*

$$\frac{dU_1}{dt} = \alpha_1 U_2 - \beta_1 P_8 U_1 \tag{S34}$$

$$\frac{dU_2}{dt} = \beta_1 P_8 U_1 - \alpha_1 U_2 \tag{S35}$$

$$0 = U_T - U_1 - U_2 \tag{S36}$$

*Transcriptional unit:*

$$\frac{dM_3}{dt} = \frac{\alpha_{3basal} + \alpha_{3\min}\left(\frac{U_1}{K_1}\right)^n + \alpha_{3\max}\left(\frac{U_2}{K_2}\right)^p}{1 + \left(\frac{U_1}{K_1}\right)^n + \left(\frac{U_2}{K_2}\right)^p} - \beta_3 M_3 \tag{S37}$$

*Protein synthesis;*

$$\frac{dT_4}{dt} = \alpha_4 M_3 - \beta_4 T_4 \tag{S38}$$

$$\frac{dGH_5}{dt} = \alpha_5 M_3 - \beta_5 GH_5 \tag{S39}$$

$$\frac{dB_6}{dt} = \alpha_6 M_3 - \beta_6 B_6 \tag{S40}$$

$$\frac{dC_7}{dt} = \alpha_7 M_3 - \beta_7 C_7 \tag{S41}$$

*Enzymatic reactions:*

$$\frac{dP_8}{dt} = \frac{T_4 k_{cat4}\left(\frac{P_0}{K_{M0}}\right)}{1+\left(\frac{P_0}{K_{M0}}\right)} - \frac{GH_5 k_{cat5}\left(\frac{P_8}{K_{M5f}}\right)\left[1-\frac{1}{K_{eq5}}\left(\frac{CM_9}{P_8}\right)\right]}{1+\left(\frac{P_8}{K_{M5f}}\right)+\left(\frac{CM_9}{M_{M5r}}\right)} - \beta_8 P_8 \quad \text{(S42)}$$

$$\begin{aligned}\frac{dCM_9}{dt} = &\frac{GH_5 k_{cat5}\left(\frac{P_8}{K_{M5f}}\right)\left[1-\frac{1}{K_{eq5}}\left(\frac{CM_9}{P_8}\right)\right]}{1+\left(\frac{P_8}{K_{M5f}}\right)+\left(\frac{CM_9}{M_{M5r}}\right)} \\ &- \frac{B_6 k_{cat6}\left(\frac{CM_9}{M_{M6f}}\right)\left[1-\frac{1}{K_{eq6}}\left(\frac{CL_{10}}{CM_9}\right)\right]}{1+\left(\frac{CM_9}{M_{M6f}}\right)+\left(\frac{CL_{10}}{K_{M6r}}\right)}\end{aligned} \quad \text{(S43)}$$

$$\frac{dCL_{10}}{dt} = \frac{B_6 k_{cat6}\left(\frac{CM_9}{M_{M6f}}\right)\left[1-\frac{1}{K_{eq6}}\left(\frac{CL_{10}}{CM_9}\right)\right]}{1+\left(\frac{CM_9}{M_{M6f}}\right)+\left(\frac{CL_{10}}{K_{M6r}}\right)} - \frac{C_7 k_{cat7}\left(\frac{CL_{10}}{K_{M7}}\right)}{1+\left(\frac{CL_{10}}{K_{M7}}\right)} \quad \text{(S44)}$$

*Pathway flux:*

$$\frac{dF}{dt} = \frac{C_7 k_{cat7}\left(\frac{CL_{10}}{K_{M7}}\right)}{1+\left(\frac{CL_{10}}{K_{M7}}\right)} - F \quad \text{(S45)}$$

*2.2 GMA Equations for the Biochemical System*

These equations can be recast into the GMA form to yield 11 differential equations, 5 algebraic constraints and 1 conservation relationship as follows:

*Differential equations:*

$$\frac{dU_1}{dt} = \alpha_1 U_2 - \beta_1 P_8 U_1 \quad \text{(S46)}$$

$$\frac{dU_2}{dt} = \beta_1 P_8 U_1 - \alpha_1 U_2 \quad \text{(S47)}$$

$$\frac{dM_3}{dt} = \alpha_{3basal} D_3^{-1} + \alpha_{3\min} K_1^{-n} U_1^n D_3^{-1} + \alpha_{3\max} K_2^{-p} U_2^p D_3^{-1} - \beta_3 M_3 \quad \text{(S48)}$$

$$\frac{dT_4}{dt} = \alpha_4 M_3 - \beta_4 T_4 \quad \text{(S49)}$$

$$\frac{dGH_5}{dt} = \alpha_5 M_3 - \beta_5 GH_5 \quad \text{(S50)}$$

$$\frac{dB_6}{dt} = \alpha_6 M_3 - \beta_6 B_6 \quad \text{(S51)}$$

$$\frac{dC_7}{dt} = \alpha_7 M_3 - \beta_7 C_7 \quad \text{(S52)}$$

$$\begin{aligned}\frac{dP_8}{dt} &= T_4 k_{cat4} K_{M0}^{-1} P_0 D_4^{-1} + GH_5 k_{cat5} K_{M5f}^{-1} K_{eq5}^{-1} CM_9 D_5^{-1} \\ &\quad - GH_5 k_{cat5} K_{M5f}^{-1} P_8 D_5^{-1} - \beta_8 P_8\end{aligned} \quad \text{(S53)}$$

$$\begin{aligned}\frac{dCM_9}{dt} &= GH_5 k_{cat5} K_{M5f}^{-1} P_8 D_5^{-1} + B_6 k_{cat6} K_{M6f}^{-1} K_{eq5}^{-1} CL_{10} D_6^{-1} \\ &\quad - GH_5 k_{cat5} K_{M5f}^{-1} K_{eq5}^{-1} CM_9 D_5^{-1} - B_6 k_{cat6} K_{M6f}^{-1} CM_9 D_6^{-1}\end{aligned} \quad \text{(S54)}$$

$$\begin{aligned}\frac{dCL_{10}}{dt} &= B_6 k_{cat6} K_{M6f}^{-1} CM_9 D_6^{-1} \\ &\quad - B_6 k_{cat6} K_{M6f}^{-1} K_{eq5}^{-1} CL_{10} D_6^{-1} - C_7 k_{cat7} K_{M7}^{-1} CL_{10} D_7^{-1}\end{aligned} \quad \text{(S55)}$$

$$\frac{dF}{dt} = C_7 k_{cat7} K_{M7}^{-1} CL_{10} D_7^{-1} - F \quad \text{(S56)}$$

*Algebraic constraints:*

$$0 = 1 + K_1^{-n} U_1^n + K_2^{-p} U_2^p - D_3 \quad \text{(S57)}$$

$$0 = 1 + K_{M0}^{-1} P_0 - D_4 \quad \text{(S58)}$$

$$0 = 1 + K_{M5f}^{-1} P_8 + K_{M5r}^{-1} CM_9 - D_5 \quad \text{(S59)}$$

$$0 = 1 + K_{M6f}^{-1} CM_9 + K_{M6r}^{-1} CL_{10} - D_6 \quad \text{(S60)}$$

$$0 = 1 + K_{M7}^{-1} CL_{10} - D_7 \quad \text{(S61)}$$

*Conservation relationship*

$$0 = U_T - U_1 - U_2 \, . \quad \text{(S62)}$$

The cooperativity (Hill number) for transcription factor binding is represented by the kinetic orders *n* and *p*. The combination of transcriptional repression and activation requires the constraints: $\alpha_{3\min} < \alpha_{3basal} < \alpha_{3\max}$. The $\alpha$ and $\beta$ parameters are rate constants for synthesis and degradation processes, respectively. The $k_{cat}$ and $K_M$ parameters used in the metabolic pathway represent turnover numbers and Michaelis constants, respectively. Auxiliary variables $D_3$ to $D_7$ replace the denominators in the biochemical kinetic expressions during the process of recasting the original ODE system into the equivalent GMA system. The 17 equations (Eqs. S46-S62) contain a total of 30 parameters. As shown in the main text, no previous knowledge (other than the constraints noted above) for any of these parameter values is required by DST3 to characterize the dynamical behavior of the system and identify a robust operating point for the system.

### Supplementary Tables

**Table S1**. *Cyclic Cases, their Signatures and Case Numbers*. Resolving the singularity contained in case 27 (see Fig. 1B) involves a Design Space analysis of the sub-system described by Eqs. S13-S16. This analysis generates six sub-cases, each one of which is generated as dictated by a *three-digit signature*. Note that the reference system of equations from which dominant terms are picked according to the case signature is the full system (Eqs. S1-S3) and not the sub-system used to resolve the singularity (Eqs. S13-S16). The case numbers for the valid sub-cases have the parent case number with an underscore followed by a number associated with the sub-case; e.g., 27_3.

| **Sub-case 27_1**<br>**[111 11 21]** | **Sub-case 27_2**<br>**[22 112 21]** | **Sub-case 27_3**<br>**[22 11 112]** |
|---|---|---|
| $\frac{dX_1}{dt} = \alpha_{11} - \beta_{11}X_1$ | $\frac{dX_1}{dt} = 2\beta_{31}X_3 - 2\beta_{12}X_1^2$ | $\frac{dX_1}{dt} = 2\beta_{31}X_3 - 2\beta_{12}X_1^2$ |
| $\frac{dX_2}{dt} = \beta_{12}X_1^2 - \beta_{23}X_2$ | $\frac{dX_2}{dt} = \frac{1}{2}\alpha 11 - \beta_{22}X_2$ | $\frac{dX_2}{dt} = \beta_{12}X_1^2 - \beta_{23}X_2$ |
| $\frac{dX_3}{dt} = \beta_{23}X_2 - \beta_{31}X_3$ | $\frac{dX_3}{dt} = \beta_{23}X_2 - \beta_{31}X_3$ | $\frac{dX_3}{dt} = \frac{1}{2}\alpha 11 - \beta_{33}X_3$ |
| **Sub-case 27_4**<br>**[311 11 21]** | **Sub-case 27_5**<br>**[22 312 21]** | **Sub-case 27_6**<br>**[22 11 312]** |
| $\frac{dX_1}{dt} = 2\alpha_{31} - \beta_{11}X_1$ | $\frac{dX_1}{dt} = 2\beta_{31}X_3 - 2\beta_{12}X_1^2$ | $\frac{dX_1}{dt} = 2\beta_{31}X_3 - 2\beta_{12}X_1^2$ |
| $\frac{dX_2}{dt} = \beta_{12}X_1^2 - \beta_{23}X_2$ | $\frac{dX_2}{dt} = \alpha_{31} - \beta_{22}X_2$ | $\frac{dX_2}{dt} = \beta_{12}X_1^2 - \beta_{23}X_2$ |
| $\frac{dX_3}{dt} = \beta_{23}X_2 - \beta_{31}X_3$ | $\frac{dX_3}{dt} = \beta_{23}X_2 - \beta_{31}X_3$ | $\frac{dX_3}{dt} = \alpha_{31} - \beta_{33}X_3$ |

**Table S2**. *Conserved Cases and their Signatures*. The synthetic network shown in Fig. 1C can be decomposed into three different cases by applying the Design Space formalism. Dominance analysis on the conservation constraint, i.e., Eq. S25, gives rise to each of these cases. Note that each case lacks $n_{cr}$ differential equations, when compared with the full system (Eqs. S22-S25). The missing differential equation in each case is identified by a pair of zeros in the case signature.

| Case 1<br>[00 11 11 11] | Case 2<br>[11 00 11 12] | Case 3<br>[11 11 00 13] |
|---|---|---|
| $\frac{dX_2}{dt} = \beta_{11}X_1 - \beta_{21}X_2$ | $\frac{dX_1}{dt} = \alpha_{11}X_3 - \beta_{11}X_1$ | $\frac{dX_1}{dt} = \alpha_{11}X_3 - \beta_{11}X_1$ |
| $\frac{dX_3}{dt} = \beta_{21}X_2 - \alpha_{11}X_3$ | $\frac{dX_3}{dt} = \beta_{21}X_2 - \alpha_{11}X_3$ | $\frac{dX_2}{dt} = \beta_{11}X_1 - \beta_{21}X_2$ |
| $0 = CR_1 - X_1$ | $0 = CR_1 - X_2$ | $0 = CR_1 - X_3$ |

**Table S3.** *Blow-Up Cases and their Case Numbers for a Single Knife-Edge Condition*. Each sub-case is defined by the violation of its knife-edge condition in one of two possible directions. Sub-case 2.1 is valid because $X_1 \to \infty$ fulfills the associated dominance condition $K_M^{-1}X_1 > 1$. The same is not true for sub-case 2.2 because $X_1 \to 0$ does not satisfy this dominance condition. Computationally determining the validity of each sub-case involves the solution of a linear program, as described by Fasani and Savageau, (2010); the only difference being the incorporation of additional inequalities to account for the violation of the knife-edge conditions and associated extreme values for the chemical pools. Since the linear program is formulated in logarithmic coordinates, a value of $\log X_1 = 12$ or $\log X_1 = -12$ is used when $X_1 \to \infty$ or $X_1 \to 0$, respectively. The case numbers for the valid sub-cases have the parent case number with a period followed by a number associated with the sub-case. The total number of sub-cases corresponds to $2^{n_{knife}}$, where $n_{knife}$ refers to the number of knife-edge conditions in the system.

| | **Sub-case 2.1** | **Sub-case 2.2** |
|---|---|---|
| **Violation of knife-edge** | $v_0 > V_M$ | $v_0 < V_M$ |
| **Implies extreme value** | $X_1 \to \infty$ | $X_1 \to 0$ |
| **Dominance condition** | $\infty \bullet K_M^{-1} > 1$ | $0 \bullet K_M^{-1} > 1$ |
| **Validity** | ***Valid*** | Not valid |

**Table S4**. *Blow-Up Cases for Two Knife-Edge Conditions*. Since $n_{knife} = 2$, there is a total of four sub-cases to analyze. Only sub-cases 2 and 3 are valid. Sub-cases 1 and 2 exhibit an inconsistent violation of their knife-edge conditions. Sub-case 1.1 for instance, dictates $\alpha_1 > \alpha_2$ and $\alpha_2 > X_1$, which implies $X_1 \to \infty$ and $X_2 \to \infty$. By replacing extreme values into these inequalities, one obtains $\alpha_2 > \infty$, which cannot be fulfilled, rendering sub-case 1.1 invalid. Dominance conditions do not exist for this example because the full system being analyzed is already an S-System.

| | **Sub-case 1.1** | **Sub-case 1.2** | **Sub-case 1.3** | **Sub-case 1.4** |
|---|---|---|---|---|
| **Violation of knife-edge** | $\alpha_1 > \alpha_2$<br>$\alpha_2 > X_1$ | $\alpha_1 < \alpha_2$<br>$\alpha_2 > X_1$ | $\alpha_1 > \alpha_2$<br>$\alpha_2 < X_1$ | $\alpha_1 < \alpha_2$<br>$\alpha_2 < X_1$ |
| **Implies extreme values** | $X_1 \to \infty$<br>$X_2 \to \infty$ | $X_1 \to 0$<br>$X_2 \to \infty$ | $X_1 \to \infty$<br>$X_2 \to 0$ | $X_1 \to 0$<br>$X_2 \to 0$ |
| **Dominance conditions** | None | None | None | None |
| **Validity** | Not valid | ***Valid*** | ***Valid*** | Not valid |

### Supplementary Figures

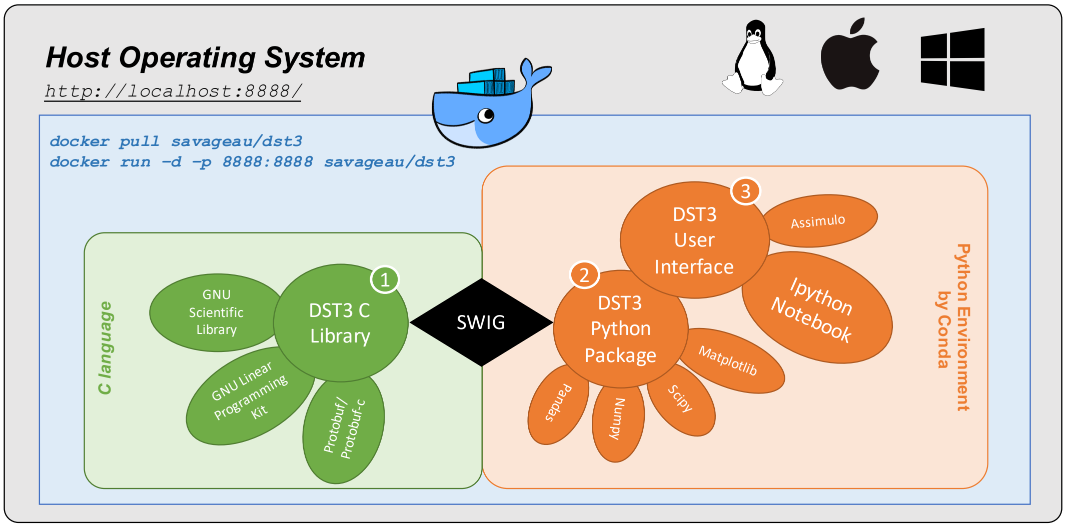


**Figure S1**. *Architecture of DST3 and Its Interaction with the Host Operating System via Docker*. The three components of DST3 are integrated in a layered fashion. The *C library* is the heart of DST3. It leverages the GNU Scientific Library (GSL) to perform numerical computations, specifically matrix operations. A customized version of the GNU Linear Programming Kit (GLPK) is used to solve linear programming problems within DST3. Google protocol buffers (Protobut/Protobuf-c) are used to write and read data. Access to the C library from Python is provided by SWIG, which stands for Simplified Wrapper and Interface Generator. This allows the creation of a *DST3 Python Package*. The *DST3 User Interface* is based on widgets provided by the IPython Notebook. The Python environment is managed by Conda. The standard Docker Image for DST3 is *savageau/dst3.* Advanced users might prefer *savageau/dst3:python3*, which comes with a DST3 Python Package for Python 3.7.3.

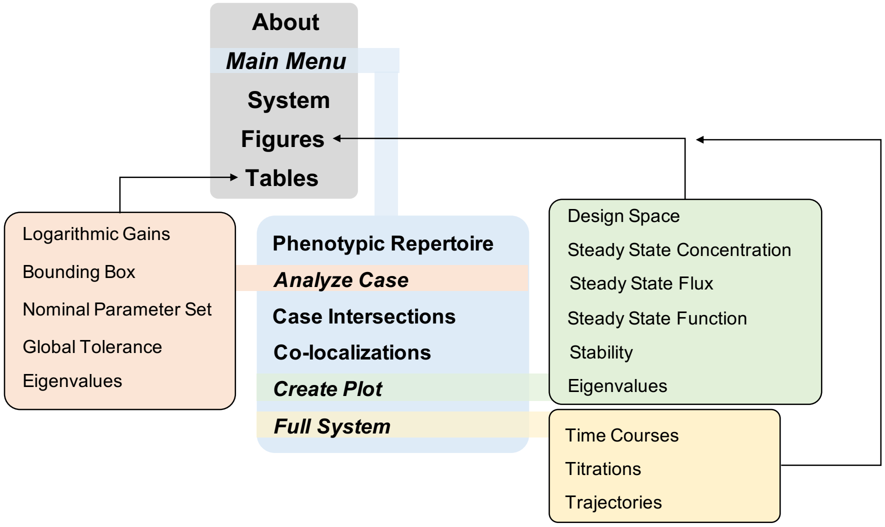


**Figure S2**. *Overview of Menus and Windows Comprising the IPython-based User Interface of DST3*. The user interface consists of a collection of tabs, buttons and text fields that facilitate the access to computational tools contained in the DST3 C library. Data can be saved to and loaded from `.dsipy` files. Tables generated from the menu *Phenotypic Repertoire* can be exported to `.xlsx` files for further analysis. Additionally, parameter values can be loaded from tables contained in files with the same extension. The User Interface of DST3 is built on legacy code inherited from DST2. Its portability is guaranteed through the virtualization technology offered by Docker.

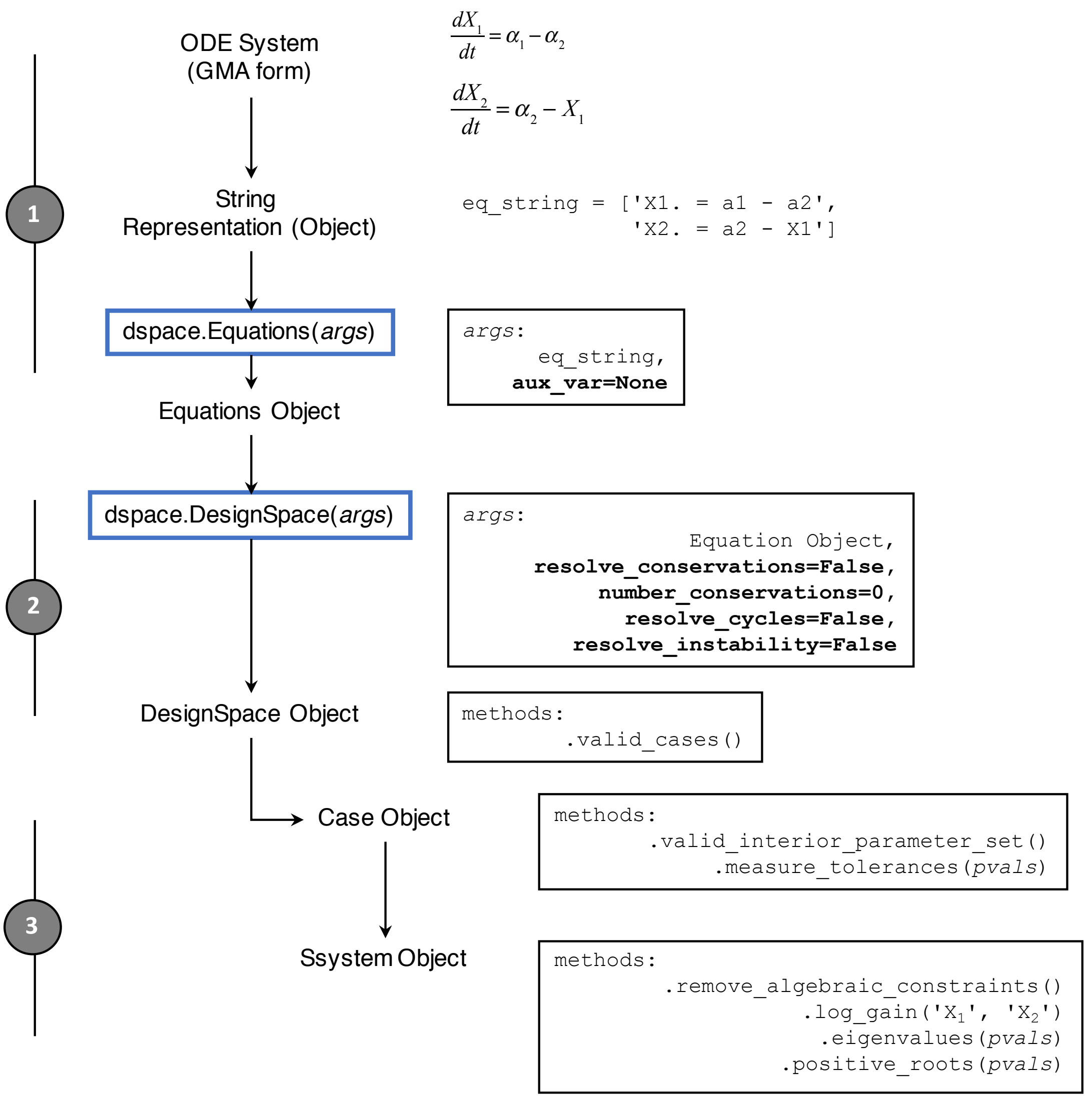


**Figure S3.** *Phenotypic Deconstruction of a Biochemical System Using the DST3 Python Module*. Three steps are involved in the computational Design Space analysis of any biochemical system. First, an `Equations` Python object is generated by means of the class `dspace.Equations`. This process involves recasting ordinary differential equations describing the systems' dynamics into a GMA form, followed by a further transformation into a list of strings according to syntax rules described in the main text. Auxiliary variables stemming from the recasting process or introduced by conservation constraints need to be declared explicitly using the key argument `aux_var`. In a second step, the `Equations` object is passed to the class `dspace.DesignSpace` along with necessary key arguments to inform the computational engine about the presence of conservations, cycles or metabolic imbalances. The output of this second step is a `DesignSpace` object. Methods associated with this object allow, among other things, the generation of a list of strings containing identifiers of valid cases. In a third step, the `DesignSpace` object can be used to generate `Case` objects using valid case identifiers as input. Each `Case` object contains a respective `SSystem` object. Methods associated with these two objects allow a comprehensive characterization of each valid case, which includes, but is by no means limited to, calculation of interior parameter sets, logarithmic gains, determination of dynamical stability, etc. Refer to Lomnitz and Savageau (2016) and to the documentation contained in the Docker Image of DST3 for more details and usage examples of the module **`dspace`**.
